# IRCAS: a novel end-to-end approach to identify, rectify and classify comprehensive alternative splicing events in a transcriptome without genome reference

**DOI:** 10.1101/2025.11.20.689457

**Authors:** Chenchen Shen, Quanbao Zhang, Qilong Cao, Xiaojun Liu, Zhen Zhang, Bailei Li, Rongqing Zhang

## Abstract

Alternative splicing (AS) is a fundamental post-transcriptional mechanism that amplifies proteomic diversity and enables adaptive responses across eukaryotes. Current AS detection methods rely heavily on reference genomes, limiting their applicability to non-model organisms. Existing reference-free approaches suffer from inaccurate splice site prediction and treat detection and classification as separate processes, resulting in cascading errors. We present IRCAS, an integrated end-to-end framework for reference-free AS analysis, comprising three modules: identification, rectification, and classification. IRCAS employs colored de Bruijn graphs for AS detection, an attention-based CNN for splice site rectification, and a hybrid Graph Neural Network combining GAT and Transformer layers for classification. Evaluation across four species demonstrates substantial improvements: splice site accuracy increased to 92-96% versus 50-55% for existing methods, and end-to-end accuracy reached 83.4% compared to 41.2% for the previous best method. IRCAS establishes a new benchmark for reference-free AS detection in non-model organisms.

**GRAPHICAL ABSTRACT:** Fig 1.
Workflow for construction and application of IRCAS. IRCAS is composed of three parts: identification, rectification, classification. (A) Workflow for reference-free AS identification from a raw transcriptomic data. First, according to the input transcripts, we apply BLAST all versus all alignment for preliminary screen. Then we adopt the MkcDBGAS Graph construction strategy. A cDBG was constructed from two sequences using a specified k-mer size. Based on bubble topologies, bubbles were classified into 5 types: SNV-induced, four AS-induced, MX-induced, AL-induced, AF-induced. (B)Workflow for AS position offset rectification and reconstruction of cDBG. Input transcript pairs are converted into a single sequence that includes two virtual nucleotides denoting the splicing start and end sites. SUPPA, a reference-based method, is utilized to determine the true splicing sites. The sequence is encoded into an n×6 vector using one-hot encoding. The offset between the true and predicted splicing sites is calculated and encoded as the ground truth. An attention-based convolutional neural network (CNN) rectification model is trained on these data to predict the offset, enabling the reconstruction of the cDBG with corrected splicing positions. (C) Workflow for 4 types AS classification. For each cDBG, node features, edge features, and global features are extracted. These features are integrated into distinct layers of a graph attention network (GAT)-Transformer hybrid model. This architecture enables high-precision classification of four types of AS events. (D)Workflow for end-to-end application of IRCAS. Transcriptomic data from any species lacking a reference genome is processed by IRCAS, enabling the classification of seven AS types with high accuracy.

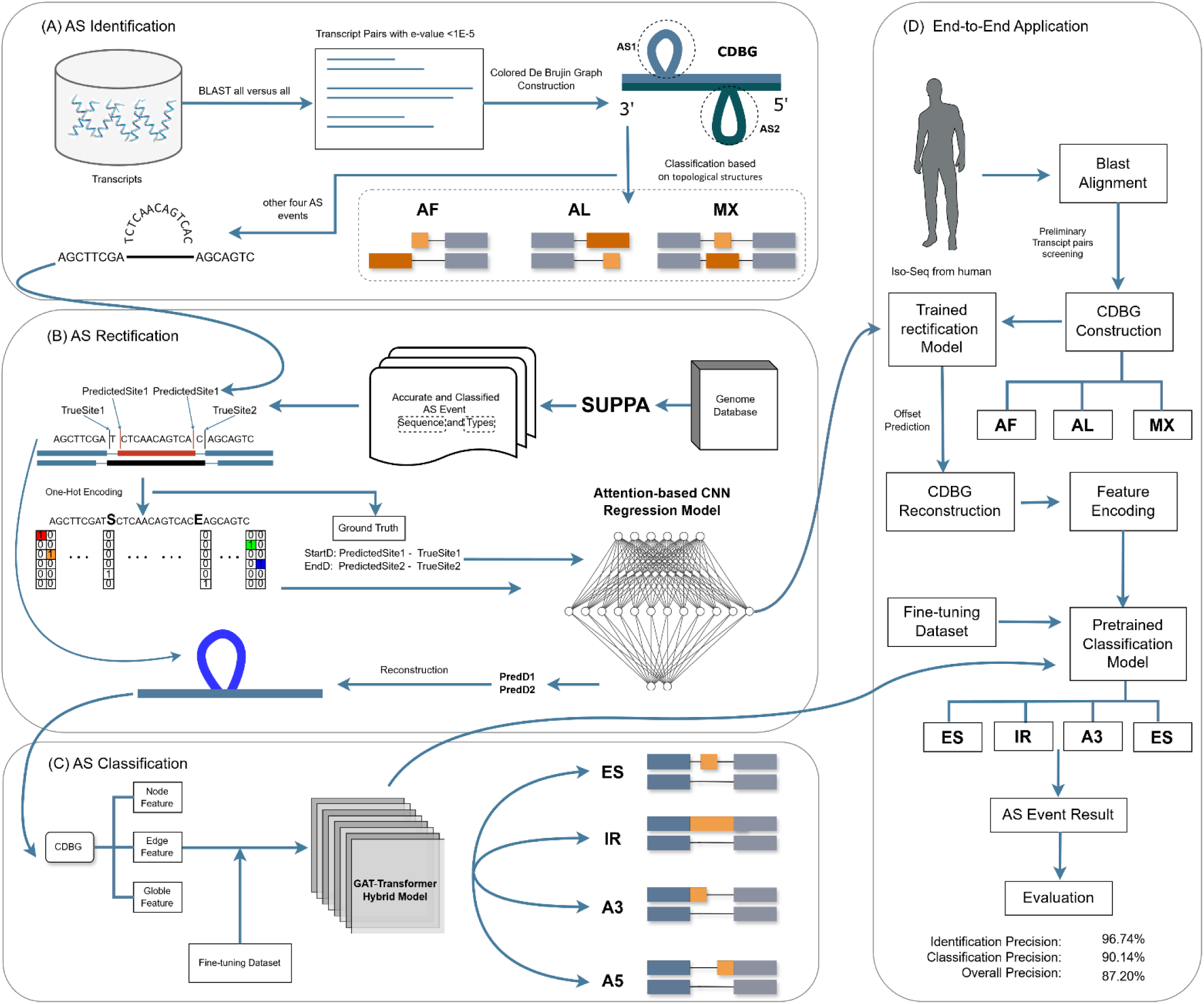

## INTRODUCTION

Alternative splicing (AS) is a post-transcriptional mechanism that increases proteomic diversity from a limited genome, contributing to phenotypic complexity and developmental plasticity in eukaryotes[1]. By enabling a single gene to generate multiple functionally distinct mRNA isoforms through differential AS events, this process fine-tunes cellular functions across tissues and developmental stages[2-4]. Critically, recent research demonstrates that AS is not merely a static feature but a dynamic modulator of environmental adaptation[5, 6]. The extensive regulatory capacity of AS establishes it as a pivotal evolutionary innovation, allowing organisms to enhance functional diversity without gene duplication and to swiftly adapt to selective pressures.[7].

RNA-Seq[8] enhances AS investigation[9], while Iso-Seq[10] enables full-length transcriptome sequencing[11]. Tools like rMATS[12], SUPPA2[13], and AStool[14] leverage these technologies to detect genome-wide AS events with corresponding genomic data. However, AS detection accuracy varies significantly, with over 70% discrepancy in differential splicing calls when using distantly related reference genomes[15, 16]. Additionally, many species lack corresponding genomic data, and acquiring such data is time-consuming and costly[17, 18]. These limitations underscore the need for reference-free AS detection methods to overcome the dependency on genomic data and improve applicability across diverse species.

Reference-free AS detection methods have evolved by improving how they identify splicing patterns in transcriptome data. Early approaches like Liu et al.[19] method utilized all-versus-all BLAST[20] to identify high-scoring segment pairs (HSPs) in order to detecting indels. These methods faced limitations in sensitivity for short splice variants, prompting AStrap[21] to integrate CD-HIT clustering with GMAP alignment[22], while IsoSplitter[23] leveraged SIM4-based alignments to detect insertion-deletion patterns characteristic of AS events, significantly improving AS detection performance. DeepASmRNA[24] enhanced the precision of AS event identification beyond AStrap by integrating biologically informed constraint. The paradigm shifted dramatically with MkcDBGAS[25], which introduced mixed k-mer colored de Bruijn graphs to dynamically identify topological "bubbles" – structures where shorter arm lengths (k-1) specifically indicate AS events. This approach achieved unprecedented precision by distinguishing paralogous genes in bubble topologies. These methodological leaps established the foundational principle that reference-free computational theory could replace reference genomes for accurate AS transcriptome pairing.

Subsequent research focused on accurate classification frameworks that transitioned from feature-dependent machine learning to end-to-end deep architectures. Seven canonical AS types are mechanistically defined: exon skipping (ES), alternative 3’ splice site (A3), alternative 5’ splice site (A5), intron retention (IR), alternative first exon (AF), alternative last exon (AL), and mutually exclusive exons (MX)[26]. AStrap pioneered AS typing with handcrafted features (splice site motifs, GC content) and tree-based classifiers, but struggled with unsatisfied precision. DeepASmRNA revolutionized this space through attention-based convolutional neural networks that directly processed one-hot encoded sequences around splice site, automatically learning discriminative patterns like "GT-AG" boundaries without manual feature engineering. The multi-scale CNN-Transformer hybrid in MCTASmRNA[27] further amplified this capability, where transformer captured long-range dependencies in sequence. Collectively, these advances represent feasible pathway toward integrated deep learning architectures capable of comprehensive AS detection and classification in reference-free contexts.

However, all previous methods have separated the prediction and classification of AS events, rather than performing an end-to-end classification starting from the raw transcript sequence. In the prediction phase, past studies evaluated the accuracy of AS event prediction by comparing correctly paired transcripts against reference-based tools like SUPPA2. While this approach provides a general assessment of prediction accuracy, it fails to precisely pinpoint the exact locations of splicing events at the base-pair level. Although predictive algorithms incorporating constrained Blast data selection, such as DeepASmRNA and MCTASmRNA, achieve over 90% accuracy in identifying correct transcript pairs, these algorithms still exhibit an average error of 2.1 bp in predicting the actual splice sites compared to their true positions. MkcDBGAS optimizes splice site selection using mixed k-mer colored de Bruijn graphs (cDBG), but it still exhibits an average position error of 1.3 bp in predicting splice sites. In the classification phase, past studies directly utilize data generated by reference-based tools which provide accurate splice sites and achieve relatively satisfied performance. In our ablation study, when using data with missing or incorrect splice sites, the model’s predictive performance significantly declined, with the previously best-performing model achieving only 74% accuracy. Critically, incorrect splice site predictions directly compromise downstream functional analysis, as even single nucleotide errors can misidentify exon-intron boundaries and lead to erroneous protein isoform annotations that fundamentally alter biological interpretations[28, 29]. In application scenarios, such positional inaccuracies not only reduce the reliability of AS event classifications but also propagate errors throughout comparative genomics studies, evolutionary analyses, and biomarker discovery workflows, resulting in a substantial loss of practical utility for researchers investigating alternative splicing mechanisms[30]. Moreover, due to the high diversity and species-specific nature of AS events, previous methods have generally failed to achieve satisfactory performance in transfer learning.

In our study, we propose IRCAS, an innovative model for the end-to-end prediction and classification of AS events, addressing the limitations of existing methodologies by seamlessly integrating accurate AS events detection with precise events classification. For raw transcripts, we employ an initial screening process utilizing BLAST to identify potential transcript pairs, complemented by the construction of cDBG to detect topological signatures indicative of AS events[31]. To enhance the accuracy of splice site localization, we introduce an attention-based convolutional neural network (CNN) regression rectification step, which refines predicted positions by leveraging contextual sequence information [15]. Subsequently, a hybrid Graph Neural Network (GNN) architecture is applied for the classification of AS event types, capitalizing on the refined positional data to achieve robust and accurate categorization. This integrated approach significantly surpasses the end-to-end accuracy of previous models, overcoming the challenges posed by inaccurate splice site predictions and enhancing the practical utility of AS detection and classification algorithms across diverse species and datasets.

## MATERIAL AND METHODS

### Data acquisition and validation set construction

To ensure comprehensive and accurate training and evaluation of the AS event detection and classification models, we selected full-length transcript datasets encompassing both animal and plant species. Specifically, human and *Arabidopsis thaliana* datasets were utilized for training, while mouse and rice datasets were employed for evaluation. All datasets and their sample sizes used in this study are detailed in Table 1. Relevant data sources are available in supplementary table S1.

**Table 1.** All AS datasets and their sample sizes.

| AS Types | Human | Arabidopsis<br>thaliana | Rice | Mouse |
| --- | --- | --- | --- | --- |
| IR | 7329 | 5992 | 2946 | 1762 |
| ES | 40 055 | 1051 | 788 | 11945 |
| A5 | 15 209 | 3161 | 1336 | 5885 |
| A3 | 16,727 | 4211 | 2472 | 7294 |
| AF | 86 273 | 936 | 235 | 6119 |
| AL | 18 923 | 106 | 61 | 3398 |
| MX | 5591 | 31 | 18 | 1526 |
| Total | 190 107 | 15 488 | 7856 | 37924 |

To train and test the IRCAS model, we applied the ‘generateEvents’ function in SUPPA2 with default settings to human and Arabidopsis thaliana annotation files, leveraging SUPPA2’s capability for accurate, systematic, and efficient genome-wide detection of all precise AS event positions and seven AS event types, which served as ground truth and labels.

### Comparing existed identification methods

To identify splice sites and clarify transcript pairs, the DeepASmRNA employed BLAST for pairwise alignment of transcript sequences with imposed constraints to detect potential splice sites. Alternatively, MkcDBGAS employs a cDBG algorithm to investigate AS events through their topological structures. Although both methods demonstrated satisfactory performance in identifying transcript pairs, they did not prioritize the accuracy of AS position detection.

### cDBG construction and AF, AL, MX classification

We employed MkcDBGAS’ workflow to identify potential alternatively spliced transcript pairs and roughly provide splicing positions. Transcript sequences were first screened for similarity using BLAST with an e-value threshold of 1e-10. For each qualifying pair, the minimum k-mer length (k) was determined as the shortest non-repetitive sequence segment to ensure an acyclic cDBG. The cDBG was then constructed as G = (V, E, C) where V comprises nodes representing k-mers from the transcripts, E includes directed edges connecting nodes with overlapping (k-1)-mers, and C denotes color labels indicating transcript origins. Bubbles[32], representing variant regions, were identified and classified topologically into single nucleotide variant (SNV)-induced (arm lengths = k), four-AS-induced (shorter arm = k-1), or other-induced (both arms > k). Three types of AS events were detected by traversing G’ to identify other-induced bubbles, applying specific criteria: a single other-induced bubble with the shorter arm <30% of the transcript length, positioned at the start (AF), end (AL), or middle (MX). For other AS types, one or more four-AS-induced bubbles with fewer than three SNV-induced bubbles were identified. The four-AS-induced category encompasses the remaining four AS event types (ES, IR, A3, A5), which exhibit identical topological structures.

### AS rectification overview

Given the substantial splicing position errors inherent in previous AS identification algorithms, we propose an attention-based convolutional neural network (CNN)[33] rectification model to address these inaccuracies. The model integrates multi-scale convolutional feature extraction with self-attention mechanisms to capture both local sequence patterns and long-range dependencies critical for splice site recognition, and outputs rectified positions with high accuracy.

### Rectification train and test set construction and data preprocessing

We used the positional offsets between cDBG-predicted splice sites and the corresponding true splice sites (determined by SUPPA2 based on the reference genome) as the ground truth for model training. Since offsets exceeding 50 bp are excessively large and extend beyond the supplemented upstream and downstream sequence boundaries, we excluded these samples during the rectification process.

The rectification model processes transcript sequences through a systematic encoding scheme that transforms nucleotide sequences into numerical representations suitable for deep learning. To address class imbalance and improve model generalization, Samples with large splice site deviations (|offset| > 3) are replicated twice. Each sequence is composed of three distinct regions: upstream sequences (50 bp), AS region (padded to uniform dimension), and downstream sequences(50 bp). Virtual nucleotides are inserted between these regions to delineate splice boundaries: ‘S’ represents the splice start site and ‘E’ represents the splice end site. The encoding strategy employs a 6-dimensional one-hot encoding for each position.

### Convolutional feature extraction and self-attention mechanism

The rectification model employs a hierarchical convolutional architecture with pogressively increasing filter complexity, consisting of four convolutional blocks: 64 filters with 3×1 kernels for local pattern detection, 128 filters with 5×1 kernels for medium-range motif recognition, 256 filters with 7×1 kernels for extended sequence context, and 512 filters with 3×1 kernels for high-level feature abstraction. Each convolutional layer is followed by batch normalization, ReLU activation, and strategic max-pooling operations (stride=2) to reduce dimensionality while preserving critical features, with dropout regularization (rate=0.1) applied to prevent overfitting. Following convolutional feature extraction, the model incorporates a multi-head self-attention module to capture long-range dependencies between distant sequence positions, comprising 8 attention heads with 512-dimensional embeddings, layer normalization applied before attention computation for training stability, and residual connections through the attention mechanism design. The attention module enables the model to dynamically focus on relevant sequence regions regardless of their positional distance, which is crucial for accurate splice site boundary detection, effectively combining the local pattern recognition capabilities of CNNs with the global context modeling of attention mechanisms.

### Huber Loss

To address the imbalanced offset distribution centered around zero, we employed Huber loss[34] to ensure robust optimization and stable convergence during training.

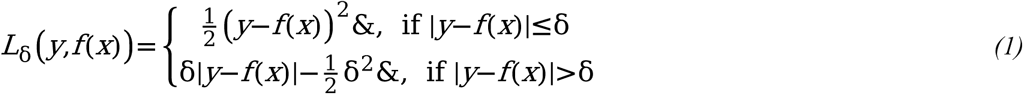

its effectiveness in addressing imbalanced offset distributions stems from its dual-regime mechanism: quadratic loss 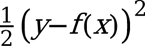 for small errors (|offset| ≤ δ) ensures precise learning of the dominant zero-centered samples, while linear loss 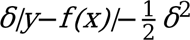 for large errors prevents rare extreme offsets from generating excessive gradients that could destabilize training. This design prevents gradient explosion from outliers while maintaining sensitivity to the majority of small offset samples, ensuring robust optimization and stable convergence.

### Reconstruction of AS events after rectification

Using the predicted offset from the rectification model, we precisely relocate splicing sites and align the upstream and downstream sequences, generating an accurate dataset for following AS classification and subsequent analyses.

### AS classification model architecture overview

To accurately classify AS events into distinct categories, we propose a hybrid graph neural network (GNN) architecture that combines Graph Attention Networks (GAT) and Transformer[35] convolution layers. The model processes cDBG graphs representing AS events, where nodes encode sequence features and edges capture splicing structure. The architecture integrates attentional aggregation mechanisms to extract discriminative graph-level representations, enabling robust classification across four AS event types despite significant class imbalance (Fig3).

**Fig 2.**
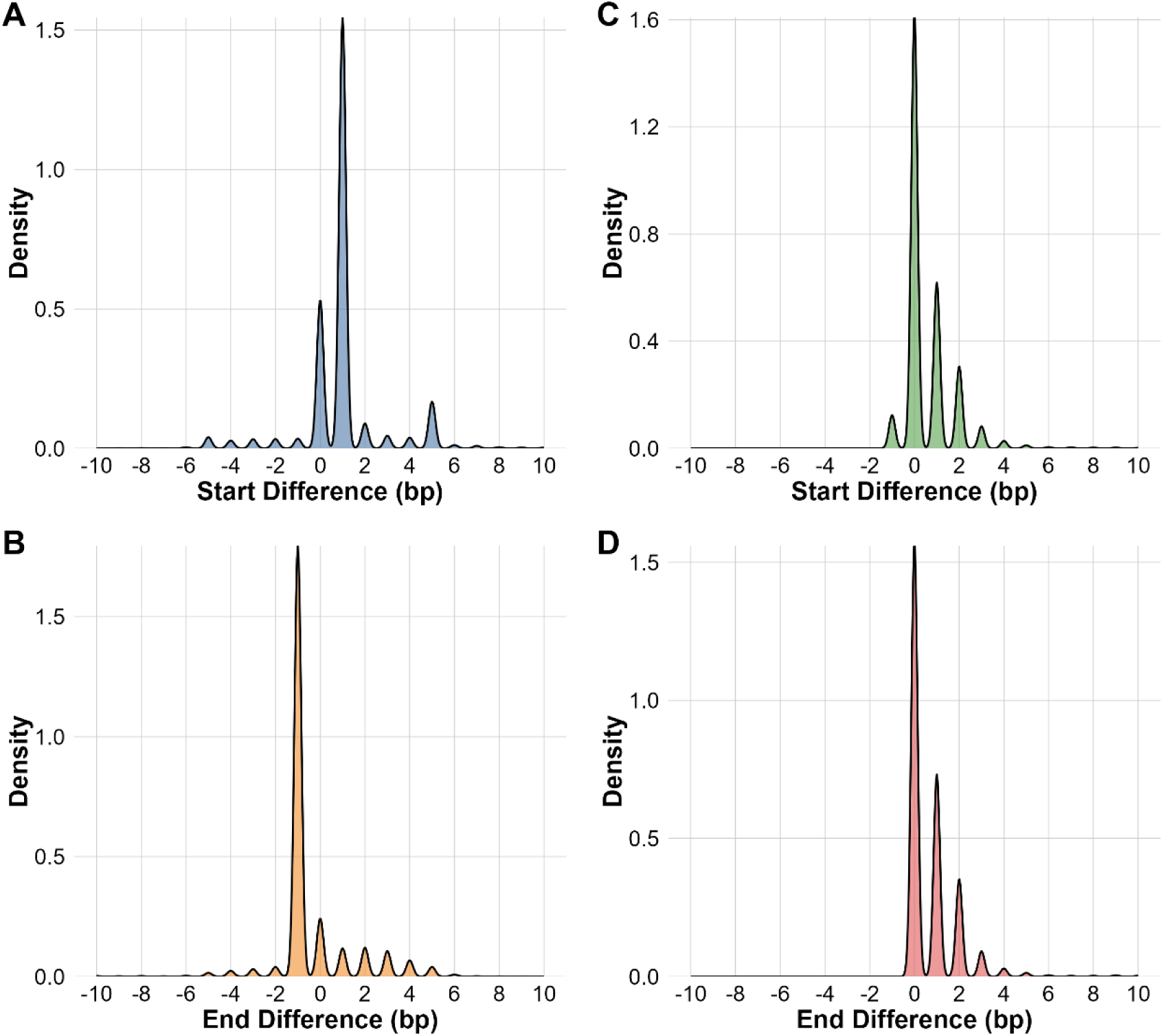
AS Position Errors Distribution by Kernel Density Estimation. (A) Distribution of splicing start position error of BLAST (B) Distribution of splicing end position error of BLAST (C) Distribution of splicing start position error of cDBG (D) Distribution of splicing end position error of cDBG

**Fig 3.**
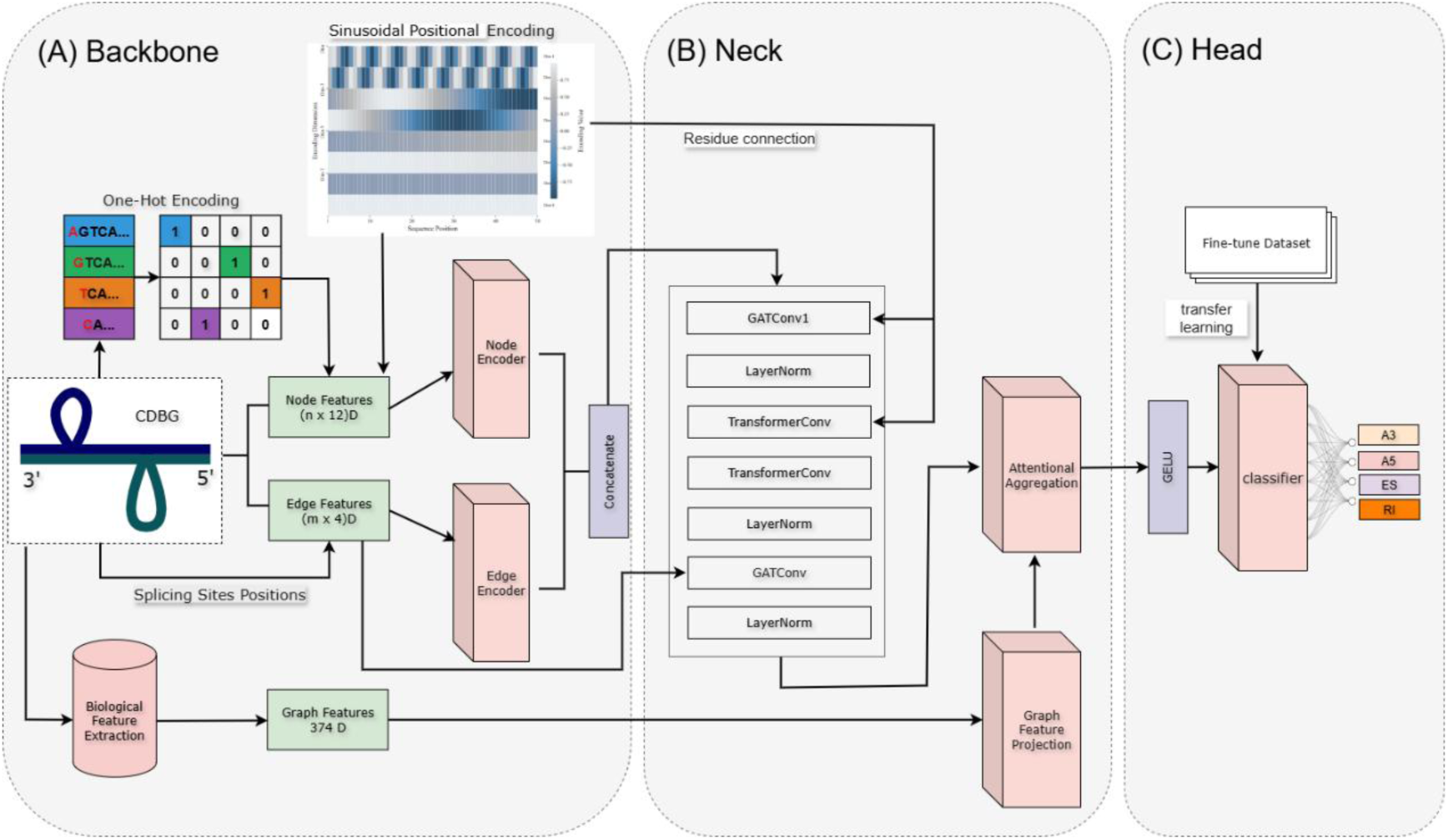
The Classification Model Architecture. (A) Backbone: The backbone encodes sequence, structural, and biological features into node, edge, and graph representations. (B) Neck: The neck employs graph neural network layers with residual connections and attentional aggregation to extract high-level graph embeddings. (C) Head: The head applying transfer learning predict AS event types.

### Classification train set construction and feature extraction

The training and test dataset comprises cDBG graph representations of AS events, with each graph labeled according to its AS type based on the result of SUPPA. We employed stratified train-validation splitting with an 80-20 ratio to preserve class distributions across datasets. Each graph contains three different levels of features: node, edge, and graph. Node features represent k-mers from the RNA sequence, encoded with a 4-dimensional one-hot vector based on the last nucleotide and an 8-dimensional sinusoidal positional encoding(PE)[36], forming a 12-dimensional feature vector.

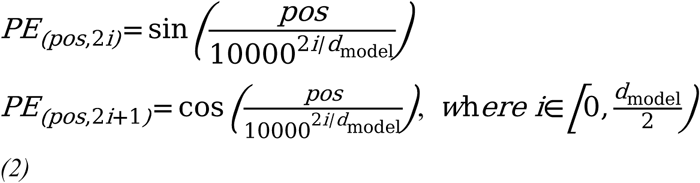

Edge features capture connections between k-mers, encoded as a 4-dimensional vector with two binary indicators for out-degree and in-degree indicating the splicing site and a 2-dimensional sinusoidal positional encoding of relative positions. Graph-level features, derived from the sequences based on validated biological features[37, 38] represented as a tensor to capture global biological properties of each AS event.

### Hybrid Graph Neural Network architecture

The classification model employs a hierarchical architecture to process cDBG graph representations for AS event classification, integrating node, edge, and graph-level features. Node features are processed through a dual-layer encoder with linear transformations, layer normalization, GELU activation, and dropout for robust feature extraction. Edge features are transformed via a dedicated edge encoder. Initially, a Graph Attention Network (GAT)[39] layer with multiple attention heads models local neighborhood interactions, followed by a projection layer. Subsequently, two Transformer layers employ edge-aware attention to capture long-range dependencies across the graph. A final single-head GAT layer refines local patterns. Each layer incorporates layer normalization, GELU activation[40], dropout, and residual connections to enhance training stability and performance. Node embeddings are augmented with positional encoding through a scaled residual connection. Graph-level representations are derived through attentional aggregation and global mean pooling, refined by a multi-head self-attention mechanism. Graph-level features are processed through a multi-layer encoder and concatenated with pooled node features. The combined representation undergoes feature fusion through a multi-layer network with linear transformations, layer normalization, GELU activation, and dropout, followed by a linear classifier producing logits for classification.

### Hybrid Loss function addressing data inbalance

To address the class imbalance prominent in the AS events dataset – where categories with fewer samples are critical – we developed a novel hybrid loss function during training. This combined cross-entropy loss [41], focal loss [42], and center loss [43] to simultaneously mitigate imbalance issues and reduce intra-class feature distances. We further enhanced this approach with dynamic weight adaptation, boosting the model’s responsiveness to all categories and overall classification accuracy.

Cross-entropy Loss: Served as the main loss function, provide stable loss for classification.

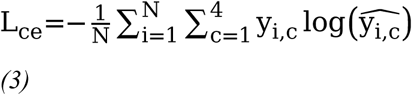

where *N* denotes the total number of samples in the dataset. The variable *C* the class. For the *i* sample, *y_i_*,*_c_* is the ground-truth label. The term 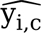 is the predicted probability that the *i* sample belongs to class *C*.

Focal Loss: Addresses class imbalance through effective number weighting and focuses learning on hard-to-classify samples.

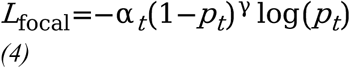

where 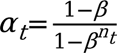 class-specific weights computed from sample frequencies n_t_ with smoothing parameter β closing to 1, and γ controls the focusing effect on misclassified examples. Additional bias factor is applied to Minority class to compensate for its severe under-representation.

Center Loss: Enhances intra-class feature compactness by minimizing feature dispersion around class centers.

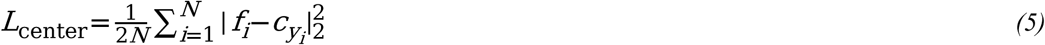

where *f_i_* denotes the feature embedding, *c_yi_* represents the learnable class center for label *y_i_*

The Hybrid loss integrates these components with adaptive normalization:

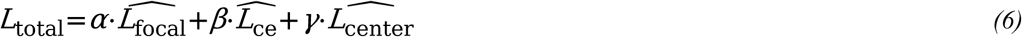

Where *α*, *β*, *γ* represents corresponding weight parameter. *α*> *β* >> *γ*.

### Evaluation and comparison to state-of-the-art model

To objectively evaluate the performance of classification model, we employed multiple metrics, including precision, recall, accuracy and F1-score for overall model assessment. To assess the efficacy of our method in classifying AS events, we benchmarked IRCAS against three state-of-the-art approaches: AStrap, DeepASmRNA, MkcDBGAS and MCTASmRNA. For AStrap, DeepASmRNA, and MkcDBGAS, we trained the models on our dataset to ensure consistency in evaluation. However, for MCTASmRNA, due to the unavailability of training code, we directly utilized the pre-trained model for testing.

### Fine-tune dataset and transfer Learning

Due to the significant variation in the characteristics and distribution of AS events across species, we employed a transfer learning strategy with a fine-tuned dataset to enhance the model’s generalization and accuracy. This approach utilized a small labeled dataset to adjust critical parameter in order to improve classification performance. In our study, we set the models trained on Human and Araport as the base model for animal and plant. For each pre-trained model, only the head layer was trained, with all other layer parameters kept frozen. To validate the significance of transfer learning and assess model generalizability, we conducted an ablation study comparing models with different dataset scales and without fine-tuning on the same dataset.

## RESULT

We introduce IRCAS, a novel tool for predicting AS events from full-length transcripts without reliance on a reference genome. IRCAS accepts full-length transcripts as input and generates 2 outputs: (i) AS transcript pairs with high credited positions of AS events, (ii) AS event types along with their confidence scores.

### Alleviating offset inaccuracy through rectification model

Previous methods for identifying AS, such as DeepASmRNA and MkcDBGAS, primarily focused on detecting transcript pairs, often neglecting precise verification of AS event positions. In the Arabidopsis dataset, these methods achieved exact AS position accuracy of only 50.1% and 53.8%, respectively. Analysis of cDBG identification splice site offsets (Table 2) revealed that approximately 56–58% of predicted start and end positions were exactly accurate, with most inaccuracies concentrated within a small range (33–39% within ±2 bp), and a minor fraction (5–6%) deviating further (up to ±10 bp or more), indicating moderate positional imprecision in prior approaches. Fig.2 illustrates the distribution of AS position identification difference density for both methods, revealing that the cDBG approach outperforms BLAST.

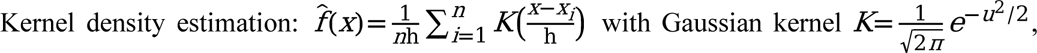

**Table 2.** cDBG identification start position and end position offset in Arabidopsis dataset.

| Interval | SoffsetCount | SoffsetPercent | EoffsetCount | EoffsetPercentage |
| --- | --- | --- | --- | --- |
| $(-\infty, -10)$ | 0 | 0.00% | 1 | 0.00% |
| $[-10, -2)$ | 0 | 0.00% | 0 | 0.00% |
| $[-2, 0)$ | 850 | 4.40% | 0 | 0.00% |
| $[0]$ | 11075 | 57.50% | 10740 | 55.80% |
| $(0, 2]$ | 6348 | 33.00% | 7443 | 38.60% |
| $(2, 10]$ | 929 | 4.80% | 1017 | 5.30% |
| $(10, \infty)$ | 59 | 0.30% | 60 | 0.30% |

where x = coordinate difference (bp), n= observation count, h= bandwidth parameter.

To evaluate the performance of our rectification models in accurately localizing AS sites across diverse species, we developed two independent models tailored to the distinct transcriptome organizations of animals and plants. The animal-trained model was developed using human data to capture splicing patterns characteristic of animal species, while the plant-trained model was trained on Arabidopsis thaliana data to account for plant-specific features. These models were designed to address lineage-specific sequence characteristics and enhance splice site prediction accuracy.

We assessed the performance of three tools—DeepASmRNA, MkcDBGAS, and our proposed IRCAS (with separate animal- and plant-trained models)—across four species: Human, Mouse, Arabidopsis, and Rice. The exact splice site accuracy (%) was measured for each model, with results summarized in a bar chart (Fig4). The animal-trained IRCAS model achieved mean accuracies of 94.3% for Human and 92.1% for Mouse, significantly outperforming DeepASmRNA (52.7% for Human, 55.2% for Mouse) and MkcDBGAS (54.1% for Human, 57.8% for Mouse). Similarly, the plant-trained IRCAS model demonstrated superior performance in plant species, with mean accuracies of 96.2% for Arabidopsis and 93.3% for Rice, compared to DeepASmRNA (43.1% for Arabidopsis, 49.6% for Rice) and MkcDBGAS (47.8% for Arabidopsis, 50.3% for Rice).

**Fig 4.**
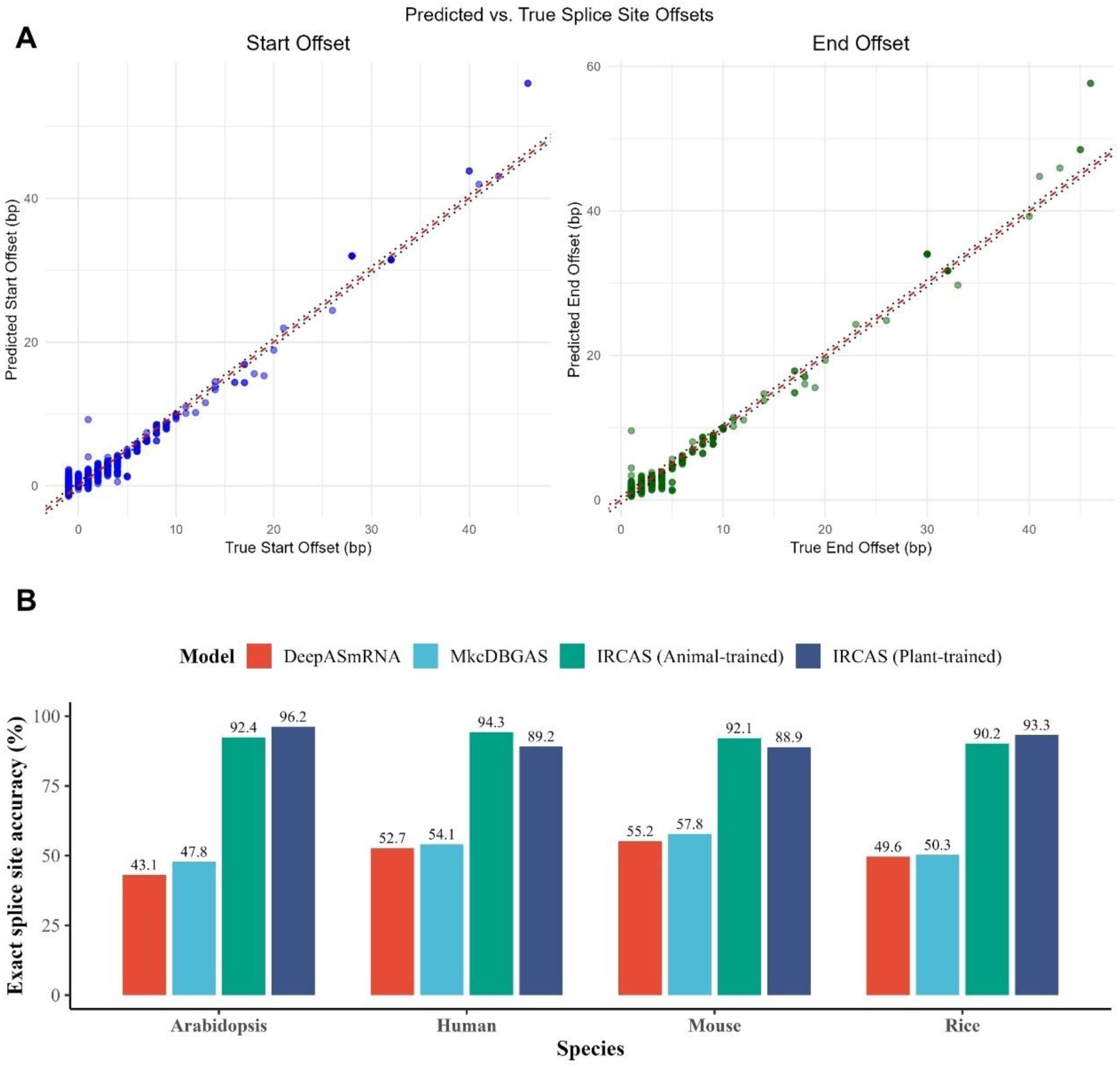
(A) Comparison of exact splice site accuracy (%) across species (Human, Mouse, Arabidopsis, Rice) for DeepASmRNA, MkcDBGAS, and IRCAS models (animal-trained and plant-trained). (B) Scatter plots of predicted versus true splice site offsets (bp) for Arabidopsis dataset with the red dashed line indicating the ideal 1:1 correlation.

The rectification step in IRCAS substantially improved splice site localization across all tested species. Compared to baseline methods, the proportion of exactly aligned splice sites increased by 34–43% across Human, Mouse, Arabidopsis, and Rice, highlighting the robustness of our attention-based convolutional neural network (CNN) rectifier. These improvements confirm that the rectification strategy effectively generalizes beyond the training organism, accommodating species-specific splicing patterns. Collectively, these results demonstrate that our species-specific training approach, combined with the rectification methodology, provides a robust and scalable framework for accurate AS event reconstruction across diverse species (Fig 4).

### A robust and highly accurate landscape of AS classification

To evaluate the effectiveness of ICRAS for AS event classification, we conducted comprehensive performance assessments across four species datasets and compared IRCAS against state-of-the-art methods. Our evaluation encompassed both individual class performance and overall classification accuracy, with particular attention to the challenging class imbalance inherent in AS event datasets.

IRCAS demonstrated superior classification performance across all tested species, achieving consistently high accuracy, precision, recall, and F1-scores. The overall classification accuracies were: Human (90.1%), Mouse (91.1%), Arabidopsis (92.8%), and Rice (91.8%).

These results represent substantial improvements over baseline methods, with particularly notable enhancements in precision and recall balance across all AS event types. The hybrid loss function effectively addressed class imbalance issues, as evidenced by the balanced performance across different AS types. Species-specific analysis revealed distinct performance patterns across taxonomic lineages. For animal species, the animal-trained model based on human dataset achieved excellent performance across both human and mouse datasets. For the human dataset, precision ranged from 83.1% (IR) to 94.4% (ES). The mouse dataset showed similar patterns, with precision values from 84.0% (IR) to 94.6% (ES). The consistent high performance across both species validates the generalizability of our animal-specific training approach. For plant species, the plant-trained model based on Arabidopsis dataset exhibited even stronger performance on plant datasets. Arabidopsis achieved the highest overall accuracy (92.8%), with precision values ranging from 89.6(ES) to 93.8(IR). The performance on rice datasets was similarly robust, exhibiting comparable trends, with precision ranging from 85.7%(ES) to 95.9% (IR). The superior performance on plant datasets may reflect the more conserved splicing patterns in plant transcriptomes compared to the complexity of animal AS. ICRAS model also Additionally, IRCAS effectively addressed minority class samples, achieving acceptable precision for ES events in plants and RI events in animals.

**Fig 5.**
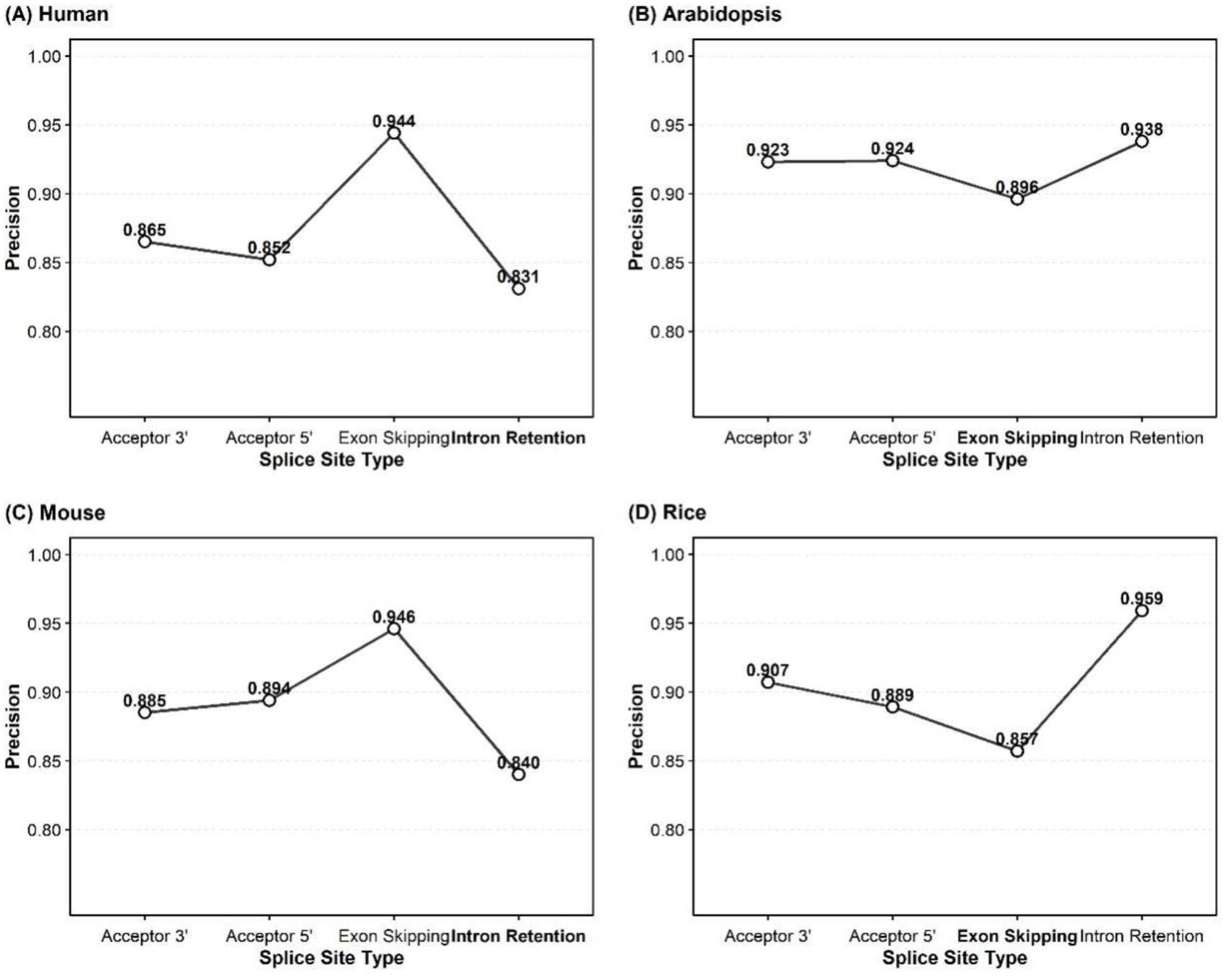
Precision of splice site prediction across four species: (A) Human, (B) Arabidopsis, (C) Mouse, and (D) Rice. Bars indicate precision values for four event types (A3, A5, ES, IR), with the least frequent splicing event type highlighted in bold on the x-axis.

### Classification performance comparing to state-of-art model

IRCAS was benchmarked against four established AS classification methods across four species (Table 4, Fig 6). IRCAS demonstrated superior classification performance across all tested species, representing consistent improvements of 1.2-4.5 percentage points over the previous best-performing method, MkcDBGAS. While DeepASmRNA and MkcDBGAS showed competitive performance (87.5-91.1%), AStrap lagged significantly with accuracies ranging from 68.6% to 84.4%, reflecting the limitations of traditional feature-based approaches compared to deep learning methods. Notably, MCTASmRNA displayed unexpectedly poor performance, with accuracies ranging from 32.4% to 55.3% across all datasets, likely attributable to the use of a pre-trained model and overfitting resulting from excessive data cleaning.

**Fig 6.**
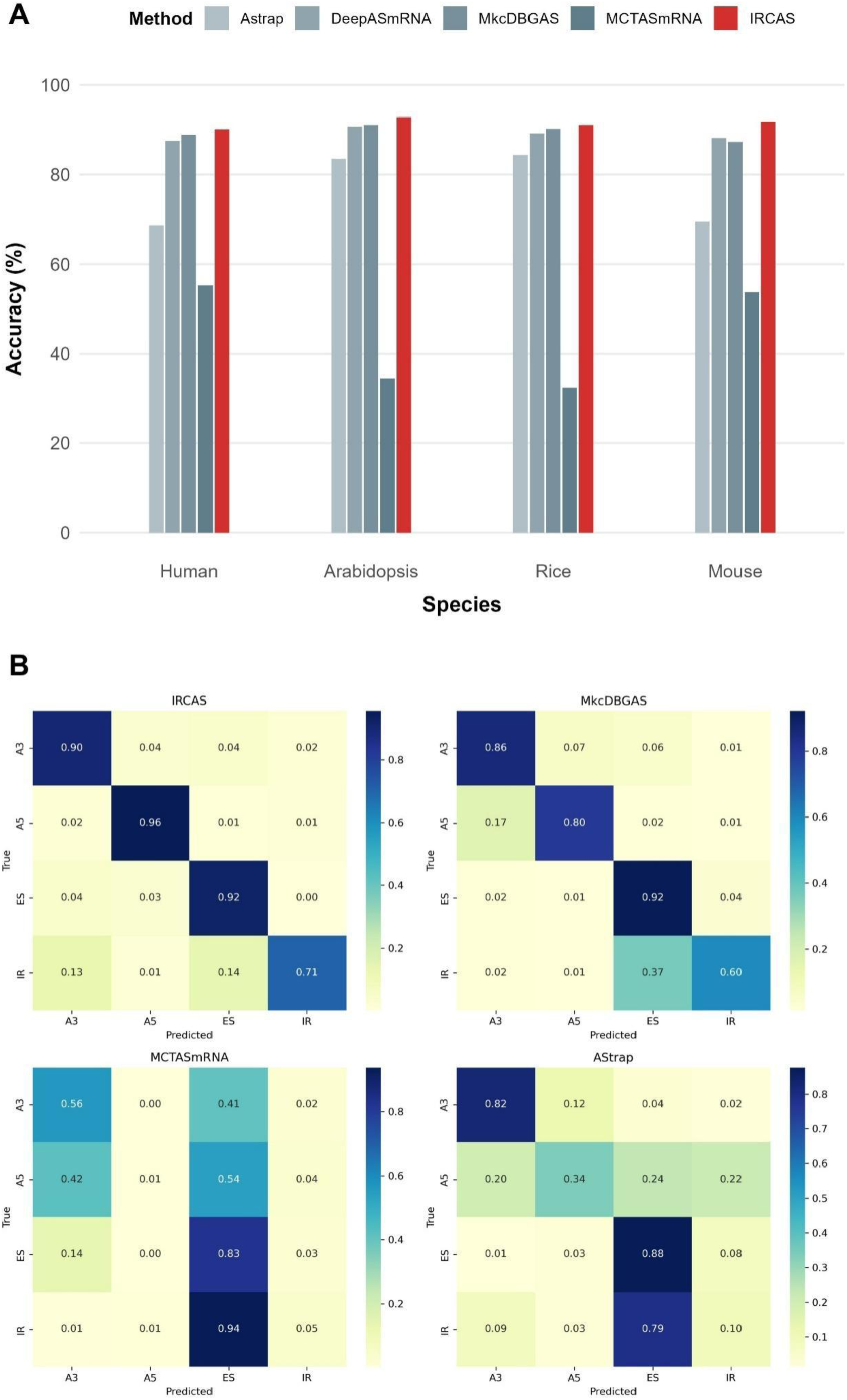
(A) Classification accuracy comparison of five AS event detection methods (AStrap, DeepASmRNA, MkcDBGAS, MCTASmRNA, IRCAS) across Human, Arabidopsis, Rice, and Mouse datasets. (B) Normalized confusion matrices heatmaps of AS event classification on human dataset across four methods (IRCAS, MkcDBGAS, MCTASmRNA, and AStrap).

**Table 3.** Prediction performance for four species was evaluated using overall accuracy, recall, F1-score, and precision for A3, A5, ES, and RI splicing events.

| Dataset | Accuracy | Recall | F1 | Precision |  |  |  |
| --- | --- | --- | --- | --- | --- | --- | --- |
|  |  |  |  | A3 | A5 | ES | IR |
| Human | 0.901 | 0.899 | 0.900 | 0.865 | 0.852 | 0.944 | 0.831 |
| Ara | 0.928 | 0.887 | 0.907 | 0.923 | 0.924 | 0.896 | 0.938 |
| Mouse | 0.911 | 0.873 | 0.892 | 0.885 | 0.894 | 0.946 | 0.840 |
| Rice | 0.918 | 0.903 | 0.910 | 0.907 | 0.889 | 0.857 | 0.959 |

**Table 4.** Overall classification accuracy (%).

| Method | Human | Arabidopsis | Rice | Mouse |
| --- | --- | --- | --- | --- |
| Astrap | 68.6 | 83.5 | 84.4 | 69.4 |
| DeepASmRNA | 87.5 | 90.7 | 89.2 | 88.2 |
| MkcDBGAS | 88.9 | 91.1 | 90.2 | 87.3 |
| MCTASmRNA | 55.3 | 34.5 | 32.4 | 53.7 |
| IRCAS | 90.1 | 92.8 | 91.9 | 91.8 |

The normalized confusion matrices heatmaps (Fig6B) further illustrate the classification patterns of the four methods. IRCAS achieved balanced and high precision across all AS event types, with relatively few off-diagonal misclassifications. In contrast, MkcDBGAS maintained competitive accuracy but exhibited higher confusion of Minority class. AStrap showed pronounced misclassifications, particularly for ES and IR events, highlighting its limited generalizability. MCTASmRNA exhibited the poorest performance, displaying widespread misclassification across categories, indicative of its instability and bias toward the majority class.

### Ablation study

To comprehensively assess the contribution of individual components within IRCAS, we conducted systematic ablation studies examining the Graph Neural Network components, the rectification module and the fine-tuning strategy across multiple performance dimensions (Fig 7).

**Fig 7.**
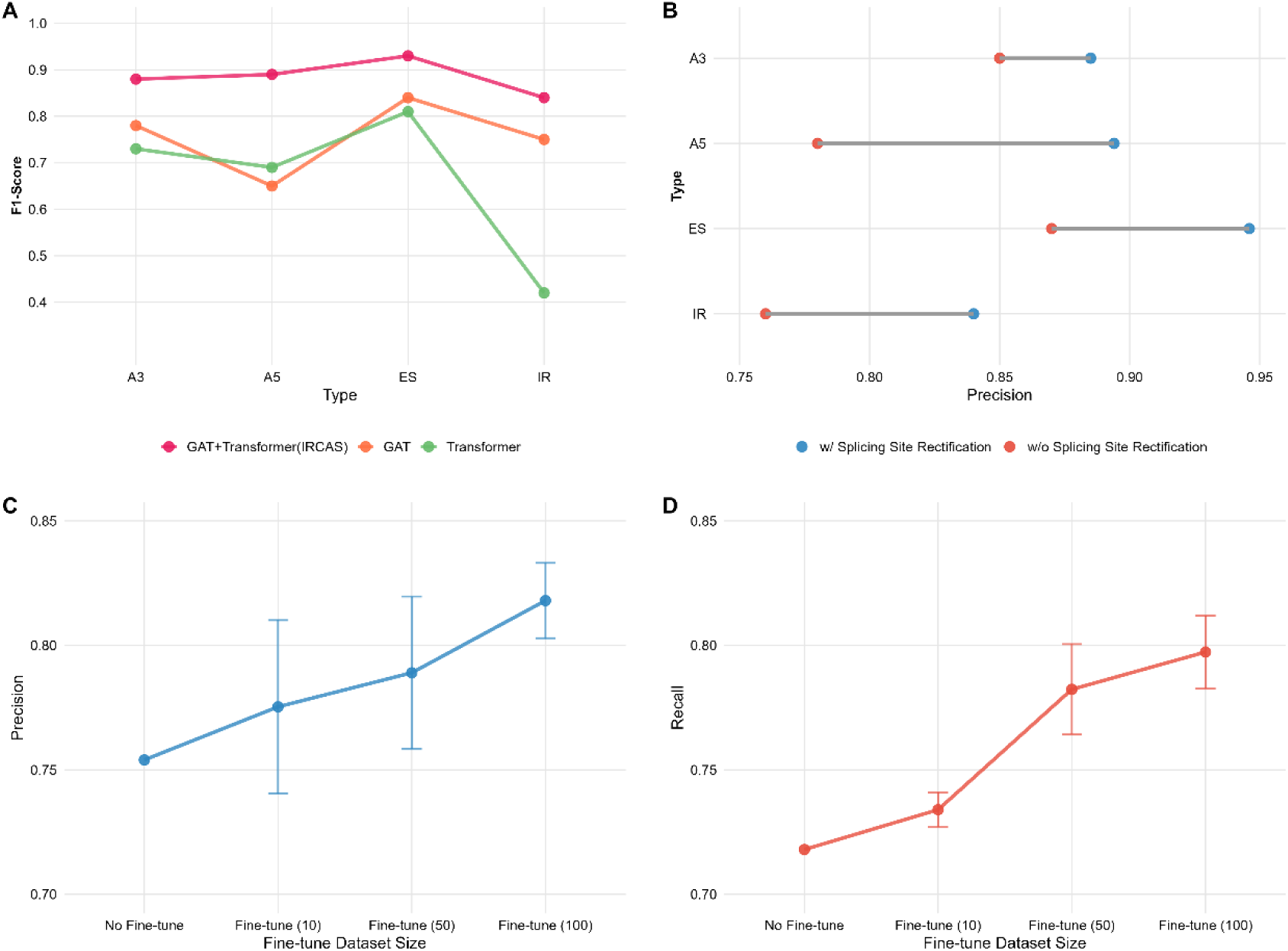
Ablation studies of IRCAS components based on mouse dataset. (A) F1-scores comparing GAT+Transformer hybrid against individual GAT and Transformer architectures. (B) Precision improvement with splicing site rectification. (C) Fine-tuning performance across dataset sizes (10, 50, 100 samples) for precision. (D) Fine-tuning performance across dataset sizes (10, 50, 100 samples) for recall. Error bars represent standard deviation from three independent random stratified sampling experiments.

To validate the effectiveness of our hybrid architecture, we compared the performance of individual components against the complete IRCAS model (Fig 6A). The GAT+Transformer(IRCAS) hybrid architecture consistently outperformed individual components across all AS event types. Compared to standalone Transformer implementation, standalone GAT implementation presents better performance, especially in Minority class (IR). These results demonstrate that the local graph relationships near the AS region captured by GAT are of paramount importance, while the Transformer’s modeling of long-range sequential patterns also significantly contributes to effective classification. The hybrid architecture successfully integrates these complementary strengths, with GAT providing structural understanding and Transformer capturing sequential dependencies that neither component adequately addresses alone.

We evaluated the impact of the splicing site rectification module in classification by comparing model performance with and without this component (Fig 6B). The rectification module demonstrated consistent improvements across all four AS event types. For A3 events, precision increased from 0.85 to 0.885. A5 events showed the most substantial improvement, rising from 0.78 to 0.894. ES events improved from 0.76 to 0.84, while IR events increased from 0.87 to 0.946. These results demonstrate that the rectification module provides crucial position refinement that significantly enhances classification accuracy across all AS types. The pronounced improvements in detecting these splice variants suggest that they are particularly sensitive to positional accuracy, likely due to subtle sequence differences that distinctly alter the features of the AS region.

To assess the relationship between fine-tuning strategy and model performance, we conducted experiments on mouse dataset using different numbers of fine-tuning samples: 10, 50, and 100 samples(Fig 6C and 6D). For precision metrics, the baseline model trained on human dataset without fine-tuning achieved 0.754. Fine-tuning with 10 samples improved performance to 0.775, representing a 2.1 percentage point increase. Scaling to 50 samples yielded further improvements to 0.789 while 100 samples achieved the highest precision of 0.818. The diminishing marginal returns observed between 50 and 100 samples suggest that substantial performance gains can be achieved with relatively small fine-tuning datasets. Similar trends were observed for recall metrics, with baseline performance at 0.718 improving to 0.734 with 10 samples, 0.782 with 50 samples, and reaching 0.797 with 100 samples. The consistent reduction in error bars with increasing sample size indicates improved model stability and generalization. Crucially, the selection of the fine-tuning dataset is pivotal to the model’s effectiveness, as non-representative datasets may compromise performance. Notably, the most substantial performance gains were observed between the baseline and 10-sample fine-tuning conditions, suggesting that even minimal fine-tuning yields significant benefits for cross-species generalization.

### Applying IRCAS to non-trained dataset and overall evaluation

To comprehensively assess IRCAS’s practical utility in reference-free AS analysis, we conducted end-to-end evaluations on the mouse dataset using models trained exclusively on human data. This cross-species evaluation framework simulates real-world applications where researchers lack reference genomes for target species but possess training data from related organisms. The overall evaluation encompasses three critical pipeline components: transcript pair identification, splicing position accuracy, and AS event type classification, with performance measured as the multiplicative product of individual component accuracies to reflect the cumulative effect of errors propagating through the pipeline.

IRCAS demonstrated superior end-to-end performance across all evaluation metrics compared to existing state-of-the-art methods. The base IRCAS model achieved 67.4% overall accuracy, representing a substantial 26.2 percentage point improvement over the previous best-performing method, MkcDBGAS (41.2%). This enhancement stems from IRCAS’s integrated approach addressing limitations in each pipeline component. While MkcDBGAS excelled in transcript pair identification (97.1%), its poor splicing position accuracy (57.8%) significantly compromised overall performance. Conversely, IRCAS maintained high transcript pair identification accuracy while dramatically improving splicing position prediction to 92.1%, demonstrating the effectiveness of our attention-based CNN rectification module.

The application of transfer learning through fine-tuning further enhanced IRCAS’s cross-species generalizability. The fine-tuned IRCAS model achieved 73.2% overall accuracy, representing a 5.8 percentage point improvement over the base model. This enhancement was primarily driven by improved AS event type classification accuracy, which increased from 75.4% to 81.8% following fine-tuning. The substantial performance gains with minimal species-specific training data (as demonstrated in our ablation studies) highlight the practical feasibility of IRCAS for non-model organisms with limited genomic resources.

Comparative analysis revealed the cumulative impact of pipeline component accuracies on overall performance. Traditional methods like AStrap suffered from poor performance across all components, achieving only 15.7% overall accuracy despite reasonable type classification performance (65.4%). DeepASmRNA showed improved transcript pair identification (90.7%) but remained limited by poor splicing position accuracy (55.2%), resulting in 36.2% overall accuracy. MCTASmRNA’s poor classification performance (52.5%) severely compromised its utility despite not requiring transcript pair identification, achieving only 26.3% overall ac These results demonstrate that IRCAS provides a robust, practical solution for reference-free AS analysis across diverse species. The model’s ability to achieve over 70% end-to-end accuracy while maintaining component-wise performance above 90% for critical tasks establishes a new benchmark for reference-free AS detection tools. The successful cross-species application from human to mouse training data, combined with the effectiveness of transfer learning approaches, positions IRCAS as a versatile tool for advancing AS research in non-model organisms where reference genomes remain unavailable.

**Table 5.** Overall application accuracy(%).

| Test Dataset | Training Dataset | Model | Transcript Pairs | Splicing Site Position | AS Events Type Classification | Overall |
| --- | --- | --- | --- | --- | --- | --- |
| Mouse | Human | AStrap | 50.8 | 47.3 | 65.4 | 15.7 |
|  |  | DeepASmRNA | 90.7 | 55.2 | 72.3 | 36.2 |
|  |  | MCTASmRNA |  |  | 52.5 | 26.3 |
|  |  | MkcDBGAS | **97.1** | 57.8 | 73.4 | 41.2 |
|  |  | IRCAS |  | **92.1** | 75.4 | 67.4 |
|  |  | IRCAS+Fine-tune |  |  | **81.8** | **73.2** |
| Rice | Arabidopsis | AStrap | 95.7 | 53.2 | 83.4 | 42.5 |
|  |  | DeepASmRNA | 94.7 | 49.6 | 87.8 | 41.2 |
|  |  | MCTASmRNA |  |  | 61.1 | 28.7 |
|  |  | MkcDBGAS | **98.7** | 50.3 | 90.1 | 44.7 |
|  |  | IRCAS |  | **93.3** | 88.7 | 81.7 |
|  |  | IRCAS+Fine-tune |  |  | **90.6** | **83.4** |

## DISCUSSION

The development of IRCAS represents a significant advancement in reference-free AS analysis by addressing fundamental limitations that have constrained previous methodologies. Previous approaches treated AS detection and classification as separate processes, creating cascading errors where inaccuracies in splice site prediction directly compromised downstream classification performance and practical application. IRCAS overcomes this limitation through an integrated end-to-end framework comprising three complementary modules: identification, rectification, and classification. The substantial improvement in splice site localization accuracy (94.3% for human, 96.2% for Arabidopsis) compared to existing methods (50-55%) addresses a critical bottleneck through our attention-based CNN rectification model, which leverages contextual sequence information to capture subtle sequence patterns that define accurate splice site.

The hybrid GNN architecture employed in IRCAS introduces several methodological innovations that collectively enhance AS classification performance. The integration of GAT and Transformer layers addresses complementary aspects of the classification problem: GAT captures local topological relationships within the cDBG structure, while Transformer layers model long-range sequence dependencies that span beyond immediate graph neighborhoods. Our ablation study demonstrates that the classification of AS events strongly relies on both local topological relationships and long-range sequence dependencies. Successful application of IRCAS across diverse species demonstrates the generalizability of learned splice site patterns and AS structural features. The species-specific training strategy, employing separate models for animal and plant, acknowledges fundamental differences in splicing mechanisms while maintaining computational efficiency. Furthermore, our transfer learning approach demonstrates that substantial performance improvements can be achieved with minimal species-specific fine-tuning data, with as few as 10 samples providing meaningful gains for cross-species adaptation.

IRCAS demonstrated superior performance across all tested species, achieving 1.2-4.5 percentage point improvements over the previous best method MkcDBGAS. The evaluation revealed distinct performance tiers: traditional feature-based approaches like AStrap showed limited effectiveness (68.6-84.4%), while deep learning methods including DeepASmRNA and MkcDBGAS achieved competitive results (87.5-91.1%). Notably, MCTASmRNA exhibited poor performance (32.4-55.3%), highlighting reproducibility challenges in deep learning AS applications. Particularly noteworthy, IRCAS demonstrated significantly higher accuracy than other models on untrained datasets, exhibiting exceptional robustness. The end-to-end accuracy of 83.4% achieved by fine-tuned IRCAS establishes a new benchmark for reference-free AS detection tools and demonstrates practical utility for non-model organisms where genome reference is limited.

Despite the significant advances achieved by IRCAS, several limitations warrant acknowledgment. First, the classification performance for animal species remains suboptimal due to the inherent complexity of animal AS events, which exhibit greater structural diversity and regulatory heterogeneity compared to plant systems. Future work could explore alternative deep learning architectures or implement species-specific sub-classification within animal lineages to better capture this complexity. Second, the computational overhead associated with constructing individual cDBG representations for each transcript pair significantly impacts training efficiency, particularly when scaling to large transcriptomic datasets. Alternative graph construction strategies or more efficient graph representation methods could address this bottleneck while maintaining the biological interpretability that makes the cDBG approach effective for AS detection.

## DATA AVAILABILITY

The code of IRCAS is available at https://github.com/S444shen/IRCAS. All relevant data is available at http://zhangqblab.cn.

## SUPPLEMENTARY DATA

Supplementary data to this article can be found at Supplementary_data.docx.

## AUTHOR CONTRIBUTIONS

## ACKNOWLEDGEMENTS

We are thankful to the editors and anonymous reviewers for insightful feedback on the manuscript.

