## supplementary data for "IRCAS: a novel end-to-end approach to identify, rectify and classify comprehensive alternative splicing events in a transcriptome without genome reference"

Table S1 Data source used by IRCAS

| Species | Data | Version | Download website |
| --- | --- | --- | --- |
| human | genome | GRCh38.p13 | https://www.gencodegenes.org/human/release_37.html |
| | annotation file | v37 | |
| | RNA-seq | PC3E | https://www.ncbi.nlm.nih.gov/sra/SRX174803 |
| | RNA-seq | GS689 | https://www.ncbi.nlm.nih.gov/sra/SRX174805 |
| mice | genome | release M31 | https://www.gencodegenes.org/mouse/release_M31.html |
| | annotation file | release M31 | |
| | transcripts | release M31 | |
| *Arabidopsis thaliana* | genome | TAIR10 | https://www.arabidopsis.org |
| | annotation file | Araport11 | |
| | transcripts | Araport11 | |
| rice | genome | Version_7.0 | https://rice.uga.edu/downloads_gad.shtml |
| | annotation file | Version_7.0 | |
| | transcripts | Version_7.0 | |

Fig S1 Training curve for rectification model.

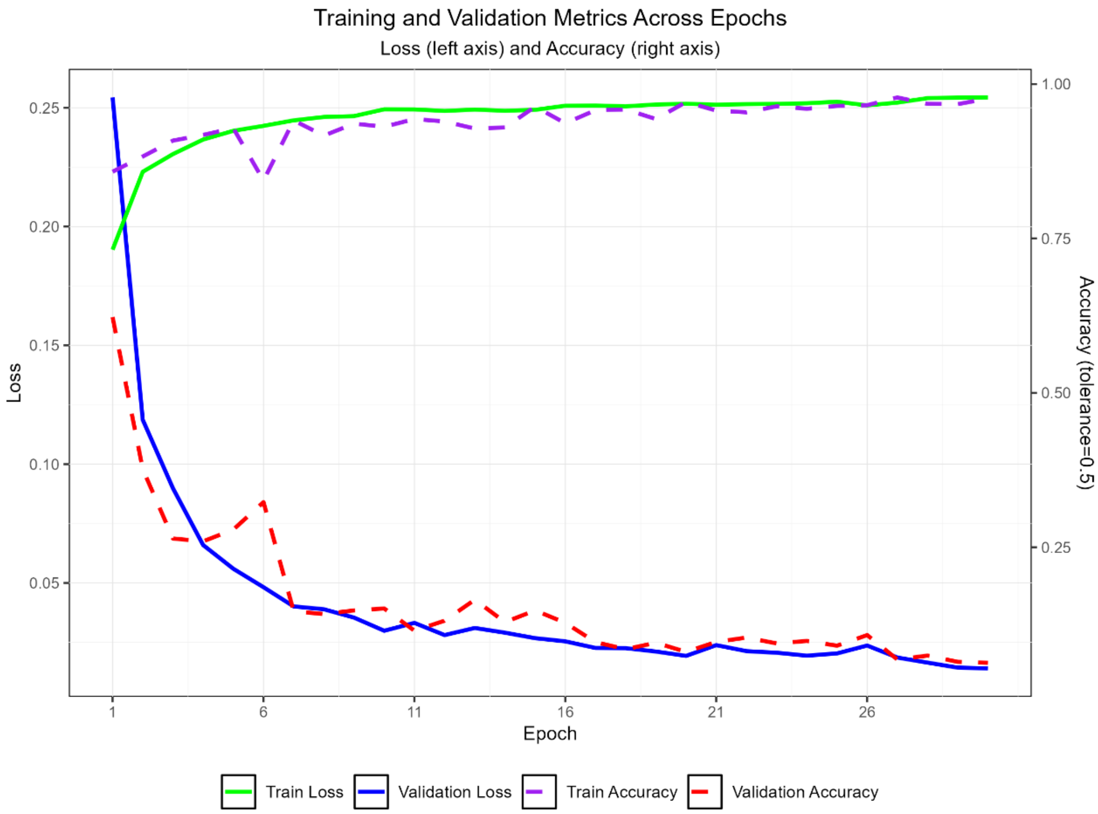

Fig S2 Training F1 curve for classification model.

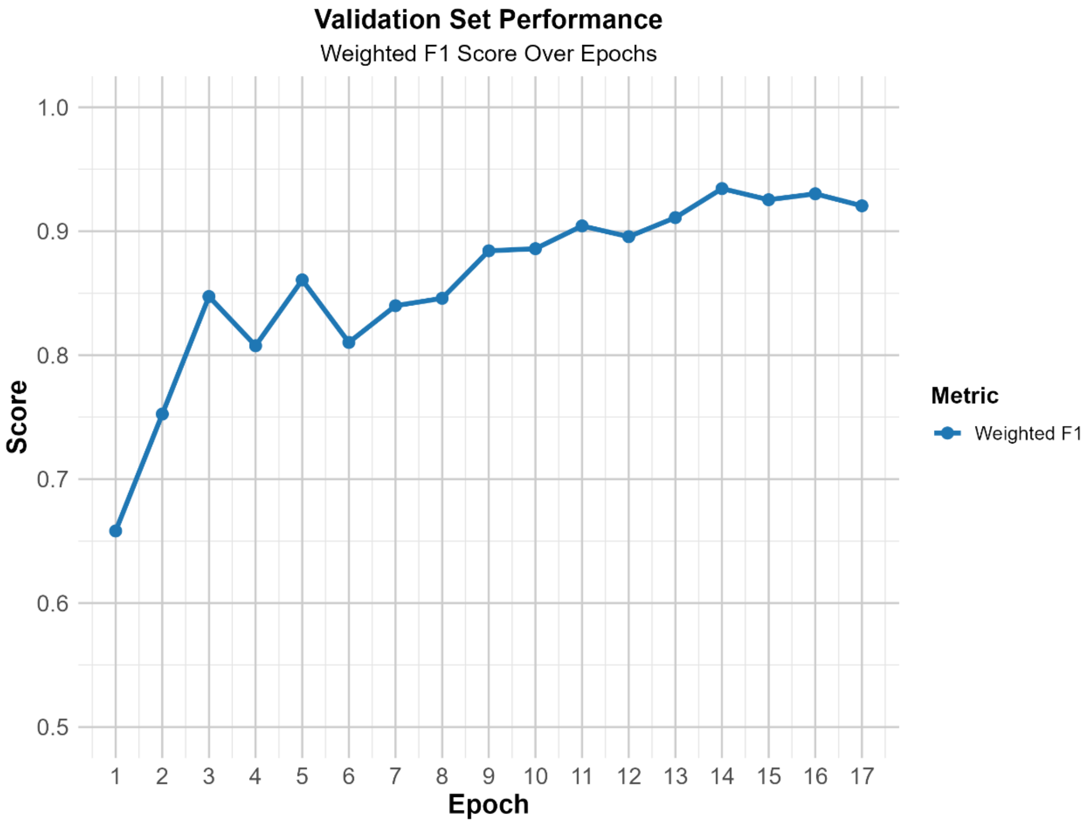

Table S2 Global features of the human model

| No. | Features | Importance | Description |
|---|---|---|---|
| 1 | length_of_as | 0.00321023 | Length of AS region |
| 2 | DmotifAAAGA | 0.00051058 | Is there this motif in downstream of splicing site: AAAGA (1 for yes, 0 for no) |
| 3 | DmotifTTCTT | 0.00061618 | Is there this motif in downstream of splicing site: TTCTT (1 for yes, 0 for no) |
| 4 | DmotifGTAGG | 0.00266274 | Is there this motif in downstream of splicing site: GTAGG (1 for yes, 0 for no) |
| 5 | DmotifTGAGG | 0.00068823 | Is there this motif in downstream of splicing site: TGAGG (1 for yes, 0 for no) |
| 6 | DmotifTCTTT | 0.00087175 | Is there this motif in downstream of splicing site: TCTTT (1 for yes, 0 for no) |
| 7 | DmotifTAAGT | 0.00069154 | Is there this motif in downstream of splicing site: TAAGT (1 for yes, 0 for no) |
| 8 | DmotifGTGAG | 0.00479803 | Is there this motif in downstream of splicing site: GTGAG (1 for yes, 0 for no) |
| 9 | DmotifGTAAG | 0.00259515 | Is there this motif in downstream of splicing site: GTAAG (1 for yes, 0 for no) |
| 10 | DmotifGTCTG | 0.00071273 | Is there this motif in downstream of splicing site: GTCTG (1 for yes, 0 for no) |
| 11 | DmotifGTTTT | 0.00067621 | Is there this motif in downstream of splicing site: GTTTT (1 for yes, |

| | | | |
|---|---|---|---|
| | | | 0 for no) |
| 12 | DmotifTTCTCT | 0.00104417 | Is there this motif in downstream of splicing site: TTCTCT (1 for yes, 0 for no) |
| 13 | DmotifTAAGG | 0.00058586 | Is there this motif in downstream of splicing site: TAAGG (1 for yes, 0 for no) |
| 14 | DmotifTCCTTT | 0.00057234 | Is there this motif in downstream of splicing site: TCCTTT (1 for yes, 0 for no) |
| 15 | DmotifTGTCT | 0.00074713 | Is there this motif in downstream of splicing site: TGTCT (1 for yes, 0 for no) |
| 16 | DmotifTTTCTC | 0.00089086 | Is there this motif in downstream of splicing site: TTTCTC (1 for yes, 0 for no) |
| 17 | DmotifCTTTT | 0.00085476 | Is there this motif in downstream of splicing site: CTTTT (1 for yes, 0 for no) |
| 18 | DmotifTGAGT | 0.00124747 | Is there this motif in downstream of splicing site: TGAGT (1 for yes, 0 for no) |
| 19 | DmotifTAGGT | 0.00133475 | Is there this motif in downstream of splicing site: TAGGT (1 for yes, 0 for no) |
| 20 | DmotifCTTTA | 0.00052679 | Is there this motif in downstream of splicing site: CTTTA (1 for yes, 0 for no) |
| 21 | DmotifTTTAG | 0.00078008 | Is there this motif in downstream of splicing site: TTTAG (1 for yes, 0 for no) |
| 22 | DmotifTGCTT | 0.00070491 | Is there this motif in downstream of splicing site: TGCTT (1 for yes, |

| | | | |
|---|---|---|---|
| | | | 0 for no) |
| 23 | DmotifGTGGGT | 0.00094222 | Is there this motif in downstream of splicing site: GTGGGT (1 for yes, 0 for no) |
| 24 | DmotifTCTCC | 0.00068126 | Is there this motif in downstream of splicing site: TCTCC (1 for yes, 0 for no) |
| 25 | DmotifTTTTTC | 0.00139376 | Is there this motif in downstream of splicing site: TTTTTC (1 for yes, 0 for no) |
| 26 | UmotifTTCTT | 0.00054549 | Is there this motif in upstream of splicing site: TTCTT (1 for yes, 0 for no) |
| 27 | UmotifTCTTT | 0.00096571 | Is there this motif in upstream of splicing site: TCTTT (1 for yes, 0 for no) |
| 28 | UmotifTAAGT | 0.00066801 | Is there this motif in upstream of splicing site: TAAGT (1 for yes, 0 for no) |
| 29 | UmotifCTCTG | 0.00061516 | Is there this motif in upstream of splicing site: CTCTG (1 for yes, 0 for no) |
| 30 | UmotifTTTTCC | 0.0007502 | Is there this motif in upstream of splicing site: TTTTCC (1 for yes, 0 for no) |
| 31 | UmotifGTGAG | 0.00550866 | Is there this motif in upstream of splicing site: GTGAG (1 for yes, 0 for no) |
| 32 | UmotifGTAAG | 0.00238317 | Is there this motif in upstream of splicing site: GTAAG (1 for yes, 0 for no) |
| 33 | UmotifCTTCT | 0.00051282 | Is there this motif in upstream of splicing site: CTTCT (1 for yes, 0 |

| | | | |
|---|---|---|---|
| | | | for no) |
| 34 | UmotifGTTTT | 0.00054749 | Is there this motif in upstream of splicing site: GTTTT (1 for yes, 0 for no) |
| 35 | UmotifCCTCT | 0.00097555 | Is there this motif in upstream of splicing site: CCTCT (1 for yes, 0 for no) |
| 36 | UmotifTCTCT | 0.00117784 | Is there this motif in upstream of splicing site: TCTCT (1 for yes, 0 for no) |
| 37 | UmotifTGTCT | 0.00059122 | Is there this motif in upstream of splicing site: TGTCT (1 for yes, 0 for no) |
| 38 | UmotifTTCCTT | 0.0005197 | Is there this motif in upstream of splicing site: TTCCTT (1 for yes, 0 for no) |
| 39 | UmotifCTTTT | 0.00080732 | Is there this motif in upstream of splicing site: CTTTT (1 for yes, 0 for no) |
| 40 | UmotifTGAGT | 0.00114328 | Is there this motif in upstream of splicing site: TGAGT (1 for yes, 0 for no) |
| 41 | UmotifCCCCAG | 0.00096588 | Is there this motif in upstream of splicing site: CCCCAG (1 for yes, 0 for no) |
| 42 | UmotifTTTAG | 0.00064961 | Is there this motif in upstream of splicing site: TTTAG (1 for yes, 0 for no) |
| 43 | UmotifTTCTC | 0.00079194 | Is there this motif in upstream of splicing site: TTCTC (1 for yes, 0 for no) |
| 44 | UmotifTGCTT | 0.00068789 | Is there this motif in upstream of splicing site: TGCTT (1 for yes, 0 |

| | | | |
|---|---|---|---|
| | | | for no) |
| 45 | allseqGC | 0.00186719 | The GC contant of all sequence |
| 46 | allseqnumberTAA | 0.00085041 | The number of stopdocon TAA in all sequence |
| 47 | allseqnumberTAG | 0.00100172 | The number of stopdocon TAG in all sequence |
| 48 | allseqnumberTGA | 0.00072766 | The number of stopdocon TGA in all sequence |
| 49 | allseqfrequencyA | 0.00094488 | The frequency of A in all sequence |
| 50 | allseqfrequencyAA | 0.00091142 | The frequency of AA in all sequence |
| 51 | allseqfrequencyAAA | 0.00050729 | The frequency of AAA in all sequence |
| 52 | allseqfrequencyAT | 0.00101279 | The frequency of AT in all sequence |
| 53 | allseqfrequencyAC | 0.00053169 | The frequency of AC in all sequence |
| 54 | allseqfrequencyACA | 0.0005611 | The frequency of ACA in all sequence |
| 55 | allseqfrequencyAGG | 0.00052174 | The frequency of AGG in all sequence |
| 56 | allseqfrequencyTT | 0.00054358 | The frequency of TT in all sequence |
| 57 | allseqfrequencyTTT | 0.00056096 | The frequency of TTT in all sequence |
| 58 | allseqfrequencyC | 0.0006169 | The frequency of C in all sequence |
| 59 | allseqfrequencyCA | 0.00096057 | The frequency of CA in all sequence |
| 60 | allseqfrequencyCAA | 0.00084891 | The frequency of CAA in all |

| | | | |
|---|---|---|---|
| | | | sequence |
| 61 | allseqfrequencyCC | 0.00064007 | The frequency of CC in all sequence |
| 62 | allseqfrequencyCCT | 0.00065494 | The frequency of CCT in all sequence |
| 63 | allseqfrequencyCCC | 0.00098559 | The frequency of CCC in all sequence |
| 64 | allseqfrequencyCCG | 0.00073578 | The frequency of CCG in all sequence |
| 65 | allseqfrequencyCG | 0.00175652 | The frequency of CG in all sequence |
| 66 | allseqfrequencyCGC | 0.00078686 | The frequency of CGC in all sequence |
| 67 | allseqfrequencyCGG | 0.00133533 | The frequency of CGG in all sequence |
| 68 | allseqfrequencyG | 0.00172969 | The frequency of G in all sequence |
| 69 | allseqfrequencyGA | 0.00059723 | The frequency of GA in all sequence |
| 70 | allseqfrequencyGAA | 0.00061731 | The frequency of GAA in all sequence |
| 71 | allseqfrequencyGTA | 0.00053189 | The frequency of GTA in all sequence |
| 72 | allseqfrequencyGC | 0.00065331 | The frequency of GC in all sequence |
| 73 | allseqfrequencyGCG | 0.00089575 | The frequency of GCG in all sequence |
| 74 | allseqfrequencyGG | 0.00104378 | The frequency of GG in all sequence |
| 75 | allseqfrequencyGGT | 0.00074696 | The frequency of GGT in all sequence |

| | | | |
|---|---|---|---|
| 76 | allseqfrequencyGGC | 0.00052334 | The frequency of GGC in all sequence |
| 77 | allseqfrequencyGGG | 0.00212446 | The frequency of GGG in all sequence |
| 78 | allseqdistr50%A | 0.00057426 | The distribution(position/length) of 50%A in all sequence |
| 79 | allseqdistr100%G | 0.00077109 | The distribution(position/length) of 100%G in all sequence |
| 80 | allseqdonerAG | 0.00066435 | Is AG in doner of all sequence (1 for yes, 0 for no) |
| 81 | asseqGC | 0.00273557 | The GC contant of AS region sequence |
| 82 | asseqnumberTAA | 0.00125697 | The number of stopdocon TAA in AS region sequence |
| 83 | asseqnumberTAG | 0.00141324 | The number of stopdocon TAG in AS region sequence |
| 84 | asseqnumberTGA | 0.0011946 | The number of stopdocon TGA in AS region sequence |
| 85 | asseqfrequencyA | 0.00209854 | The frequency of A in AS region sequence |
| 86 | asseqfrequencyAA | 0.00189235 | The frequency of AA in AS region sequence |
| 87 | asseqfrequencyAAA | 0.00068023 | The frequency of AAA in AS region sequence |
| 88 | asseqfrequencyAAT | 0.00066627 | The frequency of AAT in AS region sequence |
| 89 | asseqfrequencyAAC | 0.00065229 | The frequency of AAC in AS region sequence |
| 90 | asseqfrequencyAAG | 0.00130101 | The frequency of AAG in AS region sequence |
| 91 | asseqfrequencyAT | 0.00158739 | The frequency of AT in AS region |

| | | | |
|---|---|---|---|
| | | | sequence |
| 92 | asseqfrequencyATA | 0.00056893 | The frequency of ATA in AS region sequence |
| 93 | asseqfrequencyATT | 0.0006019 | The frequency of ATT in AS region sequence |
| 94 | asseqfrequencyATC | 0.00076242 | The frequency of ATC in AS region sequence |
| 95 | asseqfrequencyATG | 0.00077056 | The frequency of ATG in AS region sequence |
| 96 | asseqfrequencyAC | 0.00118974 | The frequency of AC in AS region sequence |
| 97 | asseqfrequencyACA | 0.00096529 | The frequency of ACA in AS region sequence |
| 98 | asseqfrequencyACT | 0.00061111 | The frequency of ACT in AS region sequence |
| 99 | asseqfrequencyACC | 0.00053821 | The frequency of ACC in AS region sequence |
| 100 | asseqfrequencyACG | 0.00055502 | The frequency of ACG in AS region sequence |
| 101 | asseqfrequencyAG | 0.0028336 | The frequency of AG in AS region sequence |
| 102 | asseqfrequencyAGA | 0.00169715 | The frequency of AGA in AS region sequence |
| 103 | asseqfrequencyAGT | 0.00094007 | The frequency of AGT in AS region sequence |
| 104 | asseqfrequencyAGC | 0.00098379 | The frequency of AGC in AS region sequence |
| 105 | asseqfrequencyAGG | 0.0015495 | The frequency of AGG in AS region sequence |
| 106 | asseqfrequencyT | 0.00200717 | The frequency of T in AS region sequence |

| | | | |
|---|---|---|---|
| 107 | asseqfrequencyTA | 0.00125172 | The frequency of TA in AS region sequence |
| 108 | asseqfrequencyTAA | 0.00063381 | The frequency of TAA in AS region sequence |
| 109 | asseqfrequencyTAT | 0.00056674 | The frequency of TAT in AS region sequence |
| 110 | asseqfrequencyTAC | 0.00060683 | The frequency of TAC in AS region sequence |
| 111 | asseqfrequencyTAG | 0.00067511 | The frequency of TAG in AS region sequence |
| 112 | asseqfrequencyTT | 0.00147658 | The frequency of TT in AS region sequence |
| 113 | asseqfrequencyTTA | 0.00055822 | The frequency of TTA in AS region sequence |
| 114 | asseqfrequencyTTT | 0.0011774 | The frequency of TTT in AS region sequence |
| 115 | asseqfrequencyTTC | 0.00084713 | The frequency of TTC in AS region sequence |
| 116 | asseqfrequencyTTG | 0.00057696 | The frequency of TTG in AS region sequence |
| 117 | asseqfrequencyTC | 0.00113531 | The frequency of TC in AS region sequence |
| 118 | asseqfrequencyTCA | 0.0012888 | The frequency of TCA in AS region sequence |
| 119 | asseqfrequencyTCT | 0.000873 | The frequency of TCT in AS region sequence |
| 120 | asseqfrequencyTCC | 0.0008606 | The frequency of TCC in AS region sequence |
| 121 | asseqfrequencyTCG | 0.00052793 | The frequency of TCG in AS region sequence |
| 122 | asseqfrequencyTG | 0.00084508 | The frequency of TG in AS region |

| | | | |
|---|---|---|---|
| | | | sequence |
| 123 | asseqfrequencyTGA | 0.00085939 | The frequency of TGA in AS region sequence |
| 124 | asseqfrequencyTGT | 0.00097791 | The frequency of TGT in AS region sequence |
| 125 | asseqfrequencyTGC | 0.00076763 | The frequency of TGC in AS region sequence |
| 126 | asseqfrequencyTGG | 0.00084692 | The frequency of TGG in AS region sequence |
| 127 | asseqfrequencyC | 0.00157183 | The frequency of C in AS region sequence |
| 128 | asseqfrequencyCA | 0.00265721 | The frequency of CA in AS region sequence |
| 129 | asseqfrequencyCAA | 0.00124177 | The frequency of CAA in AS region sequence |
| 130 | asseqfrequencyCAT | 0.00069481 | The frequency of CAT in AS region sequence |
| 131 | asseqfrequencyCAC | 0.00064456 | The frequency of CAC in AS region sequence |
| 132 | asseqfrequencyCAG | 0.00254851 | The frequency of CAG in AS region sequence |
| 133 | asseqfrequencyCT | 0.00102412 | The frequency of CT in AS region sequence |
| 134 | asseqfrequencyCTT | 0.00082232 | The frequency of CTT in AS region sequence |
| 135 | asseqfrequencyCTC | 0.00065311 | The frequency of CTC in AS region sequence |
| 136 | asseqfrequencyCTG | 0.00069029 | The frequency of CTG in AS region sequence |
| 137 | asseqfrequencyCC | 0.00125255 | The frequency of CC in AS region sequence |

| | | | |
|---|---|---|---|
| 138 | asseqfrequencyCCA | 0.00066105 | The frequency of CCA in AS region sequence |
| 139 | asseqfrequencyCCT | 0.00098925 | The frequency of CCT in AS region sequence |
| 140 | asseqfrequencyCCC | 0.00127424 | The frequency of CCC in AS region sequence |
| 141 | asseqfrequencyCCG | 0.00081053 | The frequency of CCG in AS region sequence |
| 142 | asseqfrequencyCG | 0.00158243 | The frequency of CG in AS region sequence |
| 143 | asseqfrequencyCGA | 0.00062279 | The frequency of CGA in AS region sequence |
| 144 | asseqfrequencyCGC | 0.00062939 | The frequency of CGC in AS region sequence |
| 145 | asseqfrequencyCGG | 0.00114093 | The frequency of CGG in AS region sequence |
| 146 | asseqfrequencyG | 0.00352234 | The frequency of G in AS region sequence |
| 147 | asseqfrequencyGA | 0.00239326 | The frequency of GA in AS region sequence |
| 148 | asseqfrequencyGAA | 0.00104029 | The frequency of GAA in AS region sequence |
| 149 | asseqfrequencyGAT | 0.00075181 | The frequency of GAT in AS region sequence |
| 150 | asseqfrequencyGAC | 0.00066078 | The frequency of GAC in AS region sequence |
| 151 | asseqfrequencyGAG | 0.00106982 | The frequency of GAG in AS region sequence |
| 152 | asseqfrequencyGT | 0.00298577 | The frequency of GT in AS region sequence |
| 153 | asseqfrequencyGTA | 0.00115602 | The frequency of GTA in AS |

| | | | |
|---|---|---|---|
| | | | region sequence |
| 154 | asseqfrequencyGTT | 0.00055317 | The frequency of GTT in AS region sequence |
| 155 | asseqfrequencyGTC | 0.00056694 | The frequency of GTC in AS region sequence |
| 156 | asseqfrequencyGTG | 0.00132808 | The frequency of GTG in AS region sequence |
| 157 | asseqfrequencyGC | 0.00095184 | The frequency of GC in AS region sequence |
| 158 | asseqfrequencyGCA | 0.00073113 | The frequency of GCA in AS region sequence |
| 159 | asseqfrequencyGCT | 0.00055435 | The frequency of GCT in AS region sequence |
| 160 | asseqfrequencyGCC | 0.00066351 | The frequency of GCC in AS region sequence |
| 161 | asseqfrequencyGCG | 0.00093908 | The frequency of GCG in AS region sequence |
| 162 | asseqfrequencyGG | 0.00222462 | The frequency of GG in AS region sequence |
| 163 | asseqfrequencyGGA | 0.00122948 | The frequency of GGA in AS region sequence |
| 164 | asseqfrequencyGGT | 0.00081454 | The frequency of GGT in AS region sequence |
| 165 | asseqfrequencyGGC | 0.00117968 | The frequency of GGC in AS region sequence |
| 166 | asseqfrequencyGGG | 0.00267595 | The frequency of GGG in AS region sequence |
| 167 | asseqdistr1%A | 0.00411731 | The distribution(position/length) of 1%A in AS region sequence |
| 168 | asseqdistr25%A | 0.00606579 | The distribution(position/length) of 25%A in AS region sequence |

| | | | |
|---|---|---|---|
| 169 | asseqdistr50%A | 0.00183051 | The distribution(position/length) of 50%A in AS region sequence |
| 170 | asseqdistr75%A | 0.00264277 | The distribution(position/length) of 75%A in AS region sequence |
| 171 | asseqdistr100%A | 0.00284182 | The distribution(position/length) of 100%A in AS region sequence |
| 172 | asseqdistr1%T | 0.00188463 | The distribution(position/length) of 1%T in AS region sequence |
| 173 | asseqdistr25%T | 0.00151007 | The distribution(position/length) of 25%T in AS region sequence |
| 174 | asseqdistr50%T | 0.00237935 | The distribution(position/length) of 50%T in AS region sequence |
| 175 | asseqdistr75%T | 0.00290814 | The distribution(position/length) of 75%T in AS region sequence |
| 176 | asseqdistr100%T | 0.00607481 | The distribution(position/length) of 100%T in AS region sequence |
| 177 | asseqdistr1%C | 0.00244276 | The distribution(position/length) of 1%C in AS region sequence |
| 178 | asseqdistr25%C | 0.00142548 | The distribution(position/length) of 25%C in AS region sequence |
| 179 | asseqdistr50%C | 0.00135641 | The distribution(position/length) of 50%C in AS region sequence |
| 180 | asseqdistr75%C | 0.00293903 | The distribution(position/length) of 75%C in AS region sequence |
| 181 | asseqdistr100%C | 0.00404454 | The distribution(position/length) of 100%C in AS region sequence |
| 182 | asseqdistr1%G | 0.00910389 | The distribution(position/length) of 1%G in AS region sequence |
| 183 | asseqdistr25%G | 0.00879149 | The distribution(position/length) of 25%G in AS region sequence |
| 184 | asseqdistr50%G | 0.004301 | The distribution(position/length) |

| | | | |
|---|---|---|---|
| | | | of 50%G in AS region sequence |
| 185 | asseqdistr75%G | 0.00253368 | The distribution(position/length) of 75%G in AS region sequence |
| 186 | asseqdistr100%G | 0.00106038 | The distribution(position/length) of 100%G in AS region sequence |
| 187 | asseqdonerGT | 0.00077499 | Is there GT in doner of AS region sequence (1 for yes, 0 for no) |
| 188 | asseqacceptorGT | 0.0232506 | Is there GT in acceptor of AS region sequence (1 for yes, 0 for no) |
| 189 | asseqdonerGC | 0.00053073 | Is there GC in doner of AS region sequence (1 for yes, 0 for no) |
| 190 | asseqacceptorGC | 0.00103237 | Is there GC in acceptor of AS region sequence (1 for yes, 0 for no) |
| 191 | asseqdonerAT | 0.00087271 | Is there AT in doner of AS region sequence (1 for yes, 0 for no) |
| 192 | asseqacceptorAT | 0.0011296 | Is there AT in acceptor of AS region sequence (1 for yes, 0 for no) |
| 193 | asseqdonerAG | 0.00848124 | Is there AG in doner of AS region sequence (1 for yes, 0 for no) |
| 194 | asseqacceptorAG | 0.00450759 | Is there AG in acceptor of AS region sequence (1 for yes, 0 for no) |
| 195 | asseqacceptorAC | 0.00057082 | Is there AC in acceptor of AS region sequence (1 for yes, 0 for no) |
| 196 | updistr100%A | 0.00050752 | The distribution(position/length) of 100%A in upstream sequence |
| 197 | updistr100%G | 0.00156791 | The distribution(position/length) of 100%G in upstream sequence |

| | | | |
|---|---|---|---|
| 198 | updonerAG | 0.00112393 | Is AG in doner of upstream sequence (1 for yes, 0 for no) |
| 199 | downGC | 0.00129222 | The GC contant of downstream sequence |
| 200 | downfrequencyAT | 0.00075052 | The frequency of AT in downstream sequence |
| 201 | downfrequencyT | 0.00055035 | The frequency of T in downstream sequence |
| 202 | downfrequencyCCG | 0.00058042 | The frequency of CCG in downstream sequence |
| 203 | downfrequencyCG | 0.00116404 | The frequency of CG in downstream sequence |
| 204 | downfrequencyCGC | 0.00057317 | The frequency of CGC in downstream sequence |
| 205 | downfrequencyCGG | 0.00059605 | The frequency of CGG in downstream sequence |
| 206 | downfrequencyG | 0.00052572 | The frequency of G in downstream sequence |
| 207 | downdistr100%G | 0.00063384 | The distribution(position/length) of 100%G in downstream sequence |
| 208 | downdonerAG | 0.00062139 | Is there AG in doner of downstream sequence (1 for yes, 0 for no) |
| 209 | up30as30if%3 | 0.00474754 | Whethere the length of upstream30 + AS 30bp sequence divisible by three |
| 210 | up30as30GC | 0.00097856 | The GC contant of upstream30AS 30bp + AS 30bp sequence |
| 211 | up30as30frequencyAT | 0.00059268 | The frequency of AT in upstream 30 bp + AS 30bp sequence |

| | | | |
|---|---|---|---|
| 212 | up30as30frequencyAGG | 0.00050739 | The frequency of AGG in upstream 30 bp + AS 30bp sequence |
| 213 | up30as30frequencyTAA | 0.00051832 | The frequency of TAA in upstream 30 bp + AS 30bp sequence |
| 214 | up30as30frequencyCA | 0.00054439 | The frequency of CA in upstream 30 bp + AS 30bp sequence |
| 215 | up30as30frequencyCG | 0.0005249 | The frequency of CG in upstream 30 bp + AS 30bp sequence |
| 216 | up30as30frequencyG | 0.00090055 | The frequency of G in upstream 30 bp + AS 30bp sequence |
| 217 | up30as30frequencyGT | 0.00100878 | The frequency of GT in upstream 30 bp + AS 30bp sequence |
| 218 | up30as30frequencyGTA | 0.00186856 | The frequency of GTA in upstream 30 bp + AS 30bp sequence |
| 219 | up30as30frequencyGTG | 0.00104536 | The frequency of GTG in upstream 30 bp + AS 30bp sequence |
| 220 | up30as30frequencyGG | 0.00068261 | The frequency of GG in upstream 30 bp + AS 30bp sequence |
| 221 | up30as30frequencyGGT | 0.00203348 | The frequency of GGT in upstream 30 bp + AS 30bp sequence |
| 222 | up30as30frequencyGGG | 0.00101225 | The frequency of GGG in upstream 30 bp + AS 30bp sequence |
| 223 | up30as30distr50%A | 0.00057693 | The distribution(position/length) of 50%A in upstream 30 bp + AS 30bp sequence |
| 224 | up30as30distr75%A | 0.00065496 | The distribution(position/length) of 75%A in upstream 30 bp + AS |

| | | | |
|---|---|---|---|
| | | | 30bp sequence |
| 225 | up30as30distr100%A | 0.00079364 | The distribution(position/length) of 100%A in upstream 30 bp + AS 30bp sequence |
| 226 | up30as30distr75%T | 0.00076725 | The distribution(position/length) of 75%T in upstream 30 bp + AS 30bp sequence |
| 227 | up30as30distr100%T | 0.00055656 | The distribution(position/length) of 100%T in upstream 30 bp + AS 30bp sequence |
| 228 | up30as30distr50%G | 0.00061018 | The distribution(position/length) of 50%G in upstream 30 bp + AS 30bp sequence |
| 229 | up30as30distr75%G | 0.00056901 | The distribution(position/length) of 75%G in upstream 30 bp + AS 30bp sequence |
| 230 | up30as30distr100%G | 0.00069336 | The distribution(position/length) of 100%G in upstream 30 bp + AS 30bp sequence |
| 231 | up30as30donerGT | 0.00055622 | Is there GT in doner of upstream 30 bp + AS 30bp sequence (1 for yes, 0 for no) |
| 232 | up30as30donerAG | 0.00164825 | Is there AG in doner of upstream 30 bp + AS 30bp sequence (1 for yes, 0 for no) |
| 233 | up30down30GC | 0.00052924 | The GC contant of upstream 30 + downstream 30GC sequence |
| 234 | as30down30if%3 | 0.00650359 | Whethere the length of AS 30bp + downstream 30bp sequence divisible by three |
| 235 | as30down30GC | 0.00137061 | The GC contant of AS 30bp + downstream 30bp sequence |

| | | | |
|---|---|---|---|
| 236 | as30down30frequencyA | 0.00248632 | The frequency of A in AS 30bp + downstream 30bp sequence |
| 237 | as30down30frequencyAA | 0.00118563 | The frequency of AA in AS 30bp + downstream 30bp sequence |
| 238 | as30down30frequencyAAA | 0.00069696 | The frequency of AAA in AS 30bp + downstream 30bp sequence |
| 239 | as30down30frequencyAAT | 0.00051676 | The frequency of AAT in AS 30bp + downstream 30bp sequence |
| 240 | as30down30frequencyAAG | 0.00085099 | The frequency of AAG in AS 30bp + downstream 30bp sequence |
| 241 | as30down30frequencyAT | 0.00061518 | The frequency of AT in AS 30bp + downstream 30bp sequence |
| 242 | as30down30frequencyAG | 0.0007348 | The frequency of AG in AS 30bp + downstream 30bp sequence |
| 243 | as30down30frequencyT | 0.00096321 | The frequency of T in AS 30bp + downstream 30bp sequence |
| 244 | as30down30frequencyTT | 0.00078277 | The frequency of TT in AS 30bp + downstream 30bp sequence |
| 245 | as30down30frequencyTTT | 0.00072603 | The frequency of TTT in AS 30bp + downstream 30bp sequence |
| 246 | as30down30frequencyTTC | 0.00069752 | The frequency of TTC in AS 30bp + downstream 30bp sequence |
| 247 | as30down30frequencyTC | 0.00114611 | The frequency of TC in AS 30bp + downstream 30bp sequence |
| 248 | as30down30frequencyTCT | 0.00088733 | The frequency of TCT in AS 30bp + downstream 30bp sequence |
| 249 | as30down30frequencyTCC | 0.00068028 | The frequency of TCC in AS 30bp + downstream 30bp sequence |
| 250 | as30down30frequencyTGT | 0.00050917 | The frequency of TGT in AS 30bp + downstream 30bp sequence |
| 251 | as30down30frequencyC | 0.0015108 | The frequency of C in AS 30bp + |

| | | | |
|---|---|---|---|
| | | | downstream 30bp sequence |
| 252 | as30down30frequencyCAA | 0.00057239 | The frequency of CAA in AS 30bp + downstream 30bp sequence |
| 253 | as30down30frequencyCT | 0.00100102 | The frequency of CT in AS 30bp + downstream 30bp sequence |
| 254 | as30down30frequencyCTT | 0.00062108 | The frequency of CTT in AS 30bp + downstream 30bp sequence |
| 255 | as30down30frequencyCTC | 0.00068795 | The frequency of CTC in AS 30bp + downstream 30bp sequence |
| 256 | as30down30frequencyCC | 0.00124744 | The frequency of CC in AS 30bp + downstream 30bp sequence |
| 257 | as30down30frequencyCCT | 0.00069083 | The frequency of CCT in AS 30bp + downstream 30bp sequence |
| 258 | as30down30frequencyCCC | 0.00112537 | The frequency of CCC in AS 30bp + downstream 30bp sequence |
| 259 | as30down30frequencyCG | 0.00112878 | The frequency of CG in AS 30bp + downstream 30bp sequence |
| 260 | as30down30frequencyCGC | 0.00057624 | The frequency of CGC in AS 30bp + downstream 30bp sequence |
| 261 | as30down30frequencyCGG | 0.00063691 | The frequency of CGG in AS 30bp + downstream 30bp sequence |
| 262 | as30down30frequencyG | 0.00119789 | The frequency of G in AS 30bp + downstream 30bp sequence |
| 263 | as30down30frequencyGA | 0.0007453 | The frequency of GA in AS 30bp + downstream 30bp sequence |
| 264 | as30down30frequencyGAA | 0.000782 | The frequency of GAA in AS 30bp + downstream 30bp sequence |
| 265 | as30down30frequencyGAG | 0.00055691 | The frequency of GAG in AS 30bp + downstream 30bp sequence |
| 266 | as30down30frequencyGC | 0.00059211 | The frequency of GC in AS 30bp + downstream 30bp sequence |

| | | | |
|---|---|---|---|
| 267 | as30down30frequencyGCG | 0.00084407 | The frequency of GCG in AS 30bp + downstream 30bp sequence |
| 268 | as30down30frequencyGG | 0.00070159 | The frequency of GG in AS 30bp + downstream 30bp sequence |
| 269 | as30down30frequencyGGA | 0.00054978 | The frequency of GGA in AS 30bp + downstream 30bp sequence |
| 270 | as30down30frequencyGGC | 0.0005576 | The frequency of GGC in AS 30bp + downstream 30bp sequence |
| 271 | as30down30frequencyGGG | 0.00068282 | The frequency of GGG in AS 30bp + downstream 30bp sequence |
| 272 | as30down30distr1%A | 0.00066735 | The distribution(position/length) of 1%A in AS 30bp + downstream 30bp sequence |
| 273 | as30down30distr25%A | 0.00076062 | The distribution(position/length) of 25%A in AS 30bp + downstream 30bp sequence |
| 274 | as30down30distr50%A | 0.0007277 | The distribution(position/length) of 50%A in AS 30bp + downstream 30bp sequence |
| 275 | as30down30distr75%A | 0.00051167 | The distribution(position/length) of 75%A in AS 30bp + downstream 30bp sequence |
| 276 | as30down30distr1%T | 0.00071621 | The distribution(position/length) of 1%T in AS 30bp + downstream 30bp sequence |
| 277 | as30down30distr50%T | 0.00263898 | The distribution(position/length) of 50%T in AS 30bp + downstream 30bp sequence |
| 278 | as30down30distr75%T | 0.0010547 | The distribution(position/length) of 75%T in AS 30bp + downstream 30bp sequence |
| 279 | as30down30distr1%C | 0.00056924 | The distribution(position/length) of 1%C in AS 30bp + downstream |

| | | | |
|---|---|---|---|
| | | | 30bp sequence |
| 280 | as30down30distr25%C | 0.00051178 | The distribution(position/length) of 25%C in AS 30bp + downstream 30bp sequence |
| 281 | as30down30distr50%C | 0.00052881 | The distribution(position/length) of 50%C in AS 30bp + downstream 30bp sequence |
| 282 | as30down30distr1%G | 0.00118134 | The distribution(position/length) of 1%G in AS 30bp + downstream 30bp sequence |
| 283 | as30down30distr25%G | 0.00084032 | The distribution(position/length) of 25%G in AS 30bp + downstream 30bp sequence |
| 284 | as30down30distr50%G | 0.00153808 | The distribution(position/length) of 50%G in AS 30bp + downstream 30bp sequence |
| 285 | as30down30distr75%G | 0.00058458 | The distribution(position/length) of 75%G in AS 30bp + downstream 30bp sequence |
| 286 | as30down30acceptorGT | 0.00180244 | Is there GT in acceptor of AS 30bp + downstream 30bp sequence (1 for yes, 0 for no) |
| 287 | up50as50if%3 | 0.00217258 | Whethere the length of upstream 50bp + AS 50bp sequence divisible by three |
| 288 | up50as50GC | 0.00161372 | The GC contant of upstream 50bp + AS 50bp sequence |
| 289 | up50as50numberTGA | 0.00051969 | The numberof stopdocon TGA in upstream 50bp + AS 50bp sequence |
| 290 | up50as50frequencyA | 0.00073539 | The frequency of A in upstream 50bp + AS 50bp sequence |

| | | | |
|---|---|---|---|
| 291 | up50as50frequencyAA | 0.0005253 | The frequency of AA in upstream 50bp + AS 50bp sequence |
| 292 | up50as50frequencyAAA | 0.00054986 | The frequency of AAA in upstream 50bp + AS 50bp sequence |
| 293 | up50as50frequencyAT | 0.00050509 | The frequency of AT in upstream 50bp + AS 50bp sequence |
| 294 | up50as50frequencyAGG | 0.00053668 | The frequency of AGG in upstream 50bp + AS 50bp sequence |
| 295 | up50as50frequencyCA | 0.00063607 | The frequency of CA in upstream 50bp + AS 50bp sequence |
| 296 | up50as50frequencyCC | 0.00054102 | The frequency of CC in upstream 50bp + AS 50bp sequence |
| 297 | up50as50frequencyCCT | 0.0005109 | The frequency of CCT in upstream 50bp + AS 50bp sequence |
| 298 | up50as50frequencyCCC | 0.00054381 | The frequency of CCC in upstream 50bp + AS 50bp sequence |
| 299 | up50as50frequencyCG | 0.0006102 | The frequency of CG in upstream 50bp + AS 50bp sequence |
| 300 | up50as50frequencyG | 0.0008402 | The frequency of G in upstream 50bp + AS 50bp sequence |
| 301 | up50as50frequencyGT | 0.00064936 | The frequency of GT in upstream 50bp + AS 50bp sequence |
| 302 | up50as50frequencyGTA | 0.00108651 | The frequency of GTA in upstream 50bp + AS 50bp sequence |
| 303 | up50as50frequencyGTG | 0.00062282 | The frequency of GTG in upstream 50bp + AS 50bp sequence |

| | | | |
|---|---|---|---|
| 304 | up50as50frequencyGG | 0.00088383 | The frequency of GG in upstream 50bp + AS 50bp sequence |
| 305 | up50as50frequencyGGT | 0.00109505 | The frequency of GGT in upstream 50bp + AS 50bp sequence |
| 306 | up50as50frequencyGGG | 0.00142172 | The frequency of GGG in upstream 50bp + AS 50bp sequence |
| 307 | up50as50distr25%A | 0.00052063 | The distribution(position/length) of 25%A in upstream 50bp + AS 50bp sequence |
| 308 | up50as50distr50%A | 0.00074426 | The distribution(position/length) of 50%A in upstream 50bp + AS 50bp sequence |
| 309 | up50as50distr75%A | 0.00075071 | The distribution(position/length) of 75%A in upstream 50bp + AS 50bp sequence |
| 310 | up50as50distr100%A | 0.00089748 | The distribution(position/length) of 100%A in upstream 50bp + AS 50bp sequence |
| 311 | up50as50distr50%T | 0.00061052 | The distribution(position/length) of 50%T in upstream 50bp + AS 50bp sequence |
| 312 | up50as50distr75%T | 0.00067725 | The distribution(position/length) of 75%T in upstream 50bp + AS 50bp sequence |
| 313 | up50as50distr100%T | 0.00060157 | The distribution(position/length) of 100%T in upstream 50bp + AS 50bp sequence |
| 314 | up50as50distr100%C | 0.00062694 | The distribution(position/length) of 100%C in upstream 50bp + AS 50bp sequence |
| 315 | up50as50distr25%G | 0.00051214 | The distribution(position/length) |

| | | | |
|---|---|---|---|
| | | | of 25%G in upstream 50bp + AS 50bp sequence |
| 316 | up50as50distr50%G | 0.00074642 | The distribution(position/length) of 50%G in upstream 50bp + AS 50bp sequence |
| 317 | up50as50distr75%G | 0.00067369 | The distribution(position/length) of 75%G in upstream 50bp + AS 50bp sequence |
| 318 | up50as50distr100%G | 0.0008374 | The distribution(position/length) of 100%G in upstream 50bp + AS 50bp sequence |
| 319 | up50as50donerGT | 0.00059529 | Is there GT in doner of upstream 50bp + AS 50bp sequence (1 for yes, 0 for no) |
| 320 | up50as50donerAG | 0.00252162 | Is there AG in doner of upstream 50bp + AS 50bp sequence (1 for yes, 0 for no) |
| 321 | up50down50GC | 0.00054293 | The GC contant upstream 50bp + + downstream 50bp sequence |
| 322 | up50down50distr100%G | 0.00068183 | The distribution(position/length) of 100%G in upstream 50bp + downstream 50bp sequence |
| 323 | up50down50donerAG | 0.00064568 | Is there AG in doner of upstream 50bp + downstream 50bp sequence (1 for yes, 0 for no) |
| 324 | as50down50if%3 | 0.00219184 | Whethere the length of AS 50bp + downstream 50bp sequence divisible by three |
| 325 | as50down50GC | 0.00183917 | The GC contant of GC in AS 50bp + downstream 50bp sequence |
| 326 | as50down50numberTAA | 0.00050581 | The number of stopdocon TAA in AS 50bp + downstream 50bp sequence |

| | | | |
|---|---|---|---|
| 327 | as50down50numberTGA | 0.0006772 | The number of stopdocon TGA in AS 50bp + downstream 50bp sequence |
| 328 | as50down50frequencyA | 0.00170553 | The frequency of A in AS 50bp + downstream 50bp sequence |
| 329 | as50down50frequencyAA | 0.00095836 | The frequency of AA in AS 50bp + downstream 50bp sequence |
| 330 | as50down50frequencyAAA | 0.00094627 | The frequency of AAA in AS 50bp + downstream 50bp sequence |
| 331 | as50down50frequencyAAG | 0.00079716 | The frequency of AAG in AS 50bp + downstream 50bp sequence |
| 332 | as50down50frequencyAT | 0.00081654 | The frequency of AT in AS 50bp + downstream 50bp sequence |
| 333 | as50down50frequencyATT | 0.00055137 | The frequency of ATT in AS 50bp + downstream 50bp sequence |
| 334 | as50down50frequencyAG | 0.00058994 | The frequency of AG in AS 50bp + downstream 50bp sequence |
| 335 | as50down50frequencyAGA | 0.00052927 | The frequency of AGA in AS 50bp + downstream 50bp sequence |
| 336 | as50down50frequencyT | 0.00079415 | The frequency of T in AS 50bp + downstream 50bp sequence |
| 337 | as50down50frequencyTT | 0.00059693 | The frequency of TT in AS 50bp + downstream 50bp sequence |
| 338 | as50down50frequencyTTT | 0.00068869 | The frequency of TTT in AS 50bp + downstream 50bp sequence |
| 339 | as50down50frequencyTTC | 0.00052754 | The frequency of TTC in AS 50bp + downstream 50bp sequence |
| 340 | as50down50frequencyTC | 0.00062893 | The frequency of TC in AS 50bp + downstream 50bp sequence |
| 341 | as50down50frequencyTCT | 0.00061056 | The frequency of TCT in AS 50bp + downstream 50bp sequence |

| | | | |
|---|---|---|---|
| 342 | as50down50frequencyTCC | 0.00058486 | The frequency of TCC in AS 50bp + downstream 50bp sequence |
| 343 | as50down50frequencyC | 0.00108165 | The frequency of C in AS 50bp + downstream 50bp sequence |
| 344 | as50down50frequencyCAA | 0.00084815 | The frequency of CAA in AS 50bp + downstream 50bp sequence |
| 345 | as50down50frequencyCT | 0.000901 | The frequency of CT in AS 50bp + downstream 50bp sequence |
| 346 | as50down50frequencyCTT | 0.000519 | The frequency of CTT in AS 50bp + downstream 50bp sequence |
| 347 | as50down50frequencyCTC | 0.00057286 | The frequency of CTC in AS 50bp + downstream 50bp sequence |
| 348 | as50down50frequencyCC | 0.0008822 | The frequency of CC in AS 50bp + downstream 50bp sequence |
| 349 | as50down50frequencyCCT | 0.00066117 | The frequency of CCT in AS 50bp + downstream 50bp sequence |
| 350 | as50down50frequencyCCC | 0.00123427 | The frequency of CCC in AS 50bp + downstream 50bp sequence |
| 351 | as50down50frequencyCCG | 0.00064359 | The frequency of CCG in AS 50bp + downstream 50bp sequence |
| 352 | as50down50frequencyCG | 0.00149152 | The frequency of CG in AS 50bp + downstream 50bp sequence |
| 353 | as50down50frequencyCGC | 0.00062161 | The frequency of CGC in AS 50bp + downstream 50bp sequence |
| 354 | as50down50frequencyCGG | 0.0009457 | The frequency of CGG in AS 50bp + downstream 50bp sequence |
| 355 | as50down50frequencyG | 0.00090804 | The frequency of G in AS 50bp + downstream 50bp sequence |
| 356 | as50down50frequencyGA | 0.0008211 | The frequency of GA in AS 50bp + downstream 50bp sequence |
| 357 | as50down50frequencyGAA | 0.00077229 | The frequency of GAA in AS 50bp |

| | | | |
|---|---|---|---|
| | | | + downstream 50bp sequence |
| 358 | as50down50frequencyGC | 0.00057992 | The frequency of GC in AS 50bp + downstream 50bp sequence |
| 359 | as50down50frequencyGCG | 0.00090251 | The frequency of GCG in AS 50bp + downstream 50bp sequence |
| 360 | as50down50frequencyGG | 0.00058702 | The frequency of GG in AS 50bp + downstream 50bp sequence |
| 361 | as50down50frequencyGGG | 0.00083036 | The frequency of GGG in AS 50bp + downstream 50bp sequence |
| 362 | as50down50distr1%A | 0.00076284 | The distribution(position/length) of 1%A in AS 50bp + downstream 50bp sequence |
| 363 | as50down50distr50%A | 0.00066968 | The distribution(position/length) of 50%A in AS 50bp + downstream 50bp sequence |
| 364 | as50down50distr1%T | 0.00078609 | The distribution(position/length) of 1%T in AS 50bp + downstream 50bp sequence |
| 365 | as50down50distr50%T | 0.00128706 | The distribution(position/length) of 50%T in AS 50bp + downstream 50bp sequence |
| 366 | as50down50distr75%T | 0.00079896 | The distribution(position/length) of 75%T in AS 50bp + downstream 50bp sequence |
| 367 | as50down50distr1%C | 0.00070529 | The distribution(position/length) of 1%C in AS 50bp + downstream 50bp sequence |
| 368 | as50down50distr1%G | 0.00118752 | The distribution(position/length) of 1%G in AS 50bp + downstream 50bp sequence |
| 369 | as50down50distr25%G | 0.00074848 | The distribution(position/length) of 25%G in AS 50bp + |

| | | | |
|---|---|---|---|
| | | | downstream 50bp sequence |
| 370 | as50down50distr50%G | 0.00079151 | The distribution(position/length) of 50%G in AS 50bp + downstream 50bp sequence |
| 371 | as50down50distr100%G | 0.00058932 | The distribution(position/length) of 100%G in AS 50bp + downstream 50bp sequence |
| 372 | as50down50acceptorGT | 0.00320036 | Is there GT in acceptor of AS 50bp + downstream 50bp sequence (1 for yes, 0 for no) |
| 373 | as50down50donerAG | 0.00065088 | Is there AG in doner of AS 50bp + downstream 50bp sequence (1 for yes, 0 for no) |
| 374 | as50down50acceptorAG | 0.00065056 | Is there AG in acceptor of AS 50bp + downstream 50bp sequence (1 for yes, 0 for no) |
| 375 | down20bpGC | 0.00064652 | The GC contant of downstream 20bp sequence |
| 376 | down20bpfrequencyCG | 0.00064071 | The frequency of CG in downstream 20bp sequence |

Table S3 Global features of the *Arabidopsis thaliana* model

| No. | Features | Importance | Description |
|---|---|---|---|
| 1 | length_of_as | 0.0031287 | Length of AS region |
| 2 | DmotifTTCTT | 0.00090826 | Is there this motif in downstream of splicing site: TTCTT (1 for yes, 0 for no) |
| 3 | DmotifTAACT | 0.0005252 | Is there this motif in downstream of splicing site: TAACT (1 for yes, 0 for no) |
| 4 | DmotifTCTTT | 0.00086944 | Is there this motif in downstream of splicing site: TCTTT (1 for yes, 0 for no) |
| 5 | DmotifTCTGG | 0.00050688 | Is there this motif in downstream of splicing site: TCTGG (1 for yes, 0 for no) |
| 6 | DmotifGTAAG | 0.00074236 | Is there this motif in downstream of splicing site: GTAAG (1 for yes, 0 for no) |
| 7 | DmotifGTTTT | 0.00120458 | Is there this motif in downstream of splicing site: GTTTT (1 for yes, 0 for no) |
| 8 | DmotifGTAAT | 0.00055544 | Is there this motif in downstream of splicing site: GTAAT (1 for yes, 0 for no) |
| 9 | DmotifTTCTCT | 0.00052963 | Is there this motif in downstream of splicing site: TTCTCT (1 for yes, 0 for no) |
| 10 | DmotifTATGT | 0.00129761 | Is there this motif in downstream of splicing site: TATGT (1 for yes, 0 for no) |
| 11 | DmotifTTTCTC | 0.00052936 | Is there this motif in downstream of splicing site: TTTCTC (1 for yes, 0 for no) |
| 12 | DmotifCTTTT | 0.00175439 | Is there this motif in downstream of splicing site: CTTTT (1 for yes, 0 for no) |
| 13 | DmotifTTTAG | 0.0006886 | Is there this motif in downstream of splicing site: TTTAG (1 for yes, 0 for no) |
| 14 | DmotifTTTTTC | 0.00077446 | Is there this motif in downstream of splicing site: TTTTTC (1 for yes, 0 for no) |
| 15 | DmotifTCTTG | 0.00056064 | Is there this motif in downstream of splicing site: TCTTG (1 for yes, 0 for no) |

| | | | |
|---|---|---|---|
| 16 | DmotifTCTTC | 0.00055469 | Is there this motif in downstream of splicing site: TCTTC (1 for yes, 0 for no) |
| 17 | UmotifTTCTT | 0.00099871 | Is there this motif in upstream of splicing site: TTCTT (1 for yes, 0 for no) |
| 18 | UmotifTCTTT | 0.00168493 | Is there this motif in upstream of splicing site: TCTTT (1 for yes, 0 for no) |
| 19 | UmotifCTCTG | 0.00054215 | Is there this motif in upstream of splicing site: CTCTG (1 for yes, 0 for no) |
| 20 | UmotifGTAAG | 0.00091884 | Is there this motif in upstream of splicing site: GTAAG (1 for yes, 0 for no) |
| 21 | UmotifGTTTT | 0.00129736 | Is there this motif in upstream of splicing site: GTTTT (1 for yes, 0 for no) |
| 22 | UmotifAAATT | 0.00053055 | Is there this motif in upstream of splicing site: AAATT (1 for yes, 0 for no) |
| 23 | UmotifTCTCT | 0.00069679 | Is there this motif in upstream of splicing site: TCTCT (1 for yes, 0 for no) |
| 24 | UmotifCTTTT | 0.00125088 | Is there this motif in upstream of splicing site: CTTTT (1 for yes, 0 for no) |
| 25 | UmotifTTCTC | 0.00050946 | Is there this motif in upstream of splicing site: TTCTC (1 for yes, 0 for no) |
| 26 | allseqGC | 0.00197553 | The GC contant of all sequence |
| 27 | allseqnumberTAA | 0.00093801 | The number of stopdocon TAA in all sequence |
| 28 | allseqnumberTAG | 0.00051914 | The number of stopdocon TAG in all sequence |
| 29 | allseqnumberTGA | 0.00079594 | The number of stopdocon TGA in all sequence |
| 30 | allseqfrequencyATA | 0.00054714 | The frequency of ATA in all sequence |
| 31 | allseqfrequencyAGA | 0.00056293 | The frequency of AGA in all sequence |

| | | | |
|---|---|---|---|
| 32 | allseqfrequencyT | 0.00120963 | The frequency of T in all sequence |
| 33 | allseqfrequencyTA | 0.00144838 | The frequency of TA in all sequence |
| 34 | allseqfrequencyTAA | 0.00086404 | The frequency of TAA in all sequence |
| 35 | allseqfrequencyTAT | 0.00055247 | The frequency of TAT in all sequence |
| 36 | allseqfrequencyTT | 0.00068956 | The frequency of TT in all sequence |
| 37 | allseqfrequencyTTA | 0.00065315 | The frequency of TTA in all sequence |
| 38 | allseqfrequencyTTT | 0.00102308 | The frequency of TTT in all sequence |
| 39 | allseqfrequencyCAG | 0.00080081 | The frequency of CAG in all sequence |
| 40 | allseqfrequencyCG | 0.00050537 | The frequency of CG in all sequence |
| 41 | allseqfrequencyG | 0.00064716 | The frequency of G in all sequence |
| 42 | allseqfrequencyGA | 0.00090113 | The frequency of GA in all sequence |
| 43 | allseqfrequencyGTA | 0.00082611 | The frequency of GTA in all sequence |
| 44 | allseqfrequencyGC | 0.00056062 | The frequency of GC in all sequence |
| 45 | allseqfrequencyGGA | 0.00072377 | The frequency of GGA in all sequence |
| 46 | allseqdistr1%A | 0.00050853 | The distribution(position/length) of 1% A in all sequence |
| 47 | allseqdistr25%T | 0.00075153 | The distribution(position/length) of 25% T in all sequence |
| 48 | allseqdistr50%T | 0.00061404 | The distribution(position/length) of 50% T in all sequence |
| 49 | allseqdistr75%T | 0.00062282 | The distribution(position/length) of 75% T in all sequence |
| 50 | allseqdistr25%C | 0.00053424 | The distribution(position/length) of 25% |

| | | | |
|---|---|---|---|
| | | | C in all sequence |
| 51 | allseqdistr50%C | 0.00078249 | The distribution(position/length) of 50% C in all sequence |
| 52 | allseqdistr1%G | 0.00053172 | The distribution(position/length) of 1% G in all sequence |
| 53 | allseqdistr25%G | 0.00054524 | The distribution(position/length) of 25% G in all sequence |
| 54 | allseqdistr50%G | 0.00058275 | The distribution(position/length) of 50% G in all sequence |
| 55 | allseqdistr75%G | 0.00100307 | The distribution(position/length) of 75% G in all sequence |
| 56 | allseqdistr100%G | 0.00058178 | The distribution(position/length) of 100% G in all sequence |
| 57 | asseqif%3 | 0.00075603 | Whethere the length of AS region sequence divisible by three |
| 58 | asseqGC | 0.00267662 | The GC contant of AS region sequence |
| 59 | asseqnumberTAA | 0.00154151 | The number of stopdocon TAA in AS region sequence |
| 60 | asseqnumberTAG | 0.00105079 | The number of stopdocon TAG in AS region sequence |
| 61 | asseqnumberTGA | 0.0013909 | The number of stopdocon TGA in AS region sequence |
| 62 | asseqfrequencyA | 0.0013278 | The frequency of A in AS region sequence |
| 63 | asseqfrequencyAA | 0.00189244 | The frequency of AA in AS region sequence |
| 64 | asseqfrequencyAAA | 0.00073439 | The frequency of AAA in AS region sequence |
| 65 | asseqfrequencyAAT | 0.0006585 | The frequency of AAT in AS region sequence |
| 66 | asseqfrequencyAA | 0.00072175 | The frequency of AAC in AS region |

| | | | |
|---|---|---|---|
| | C | | sequence |
| 67 | asseqfrequencyAAG | 0.00102058 | The frequency of AAG in AS region sequence |
| 68 | asseqfrequencyAT | 0.00071711 | The frequency of AT in AS region sequence |
| 69 | asseqfrequencyATA | 0.00074491 | The frequency of ATA in AS region sequence |
| 70 | asseqfrequencyATT | 0.00098504 | The frequency of ATT in AS region sequence |
| 71 | asseqfrequencyATC | 0.0007468 | The frequency of ATC in AS region sequence |
| 72 | asseqfrequencyATG | 0.00079964 | The frequency of ATG in AS region sequence |
| 73 | asseqfrequencyAC | 0.00132906 | The frequency of AC in AS region sequence |
| 74 | asseqfrequencyACA | 0.00084261 | The frequency of ACA in AS region sequence |
| 75 | asseqfrequencyACT | 0.00096637 | The frequency of ACT in AS region sequence |
| 76 | asseqfrequencyACG | 0.0005231 | The frequency of ACG in AS region sequence |
| 77 | asseqfrequencyAG | 0.00310413 | The frequency of AG in AS region sequence |
| 78 | asseqfrequencyAGA | 0.00105 | The frequency of AGA in AS region sequence |
| 79 | asseqfrequencyAGT | 0.00071315 | The frequency of AGT in AS region sequence |
| 80 | asseqfrequencyAGC | 0.00061105 | The frequency of AGC in AS region sequence |
| 81 | asseqfrequencyAGG | 0.00095123 | The frequency of AGG in AS region sequence |

| | | | |
|---|---|---|---|
| 82 | asseqfrequencyT | 0.0015526 | The frequency of T in AS region sequence |
| 83 | asseqfrequencyTA | 0.00210749 | The frequency of TA in AS region sequence |
| 84 | asseqfrequencyTA A | 0.00279647 | The frequency of TAA in AS region sequence |
| 85 | asseqfrequencyTA T | 0.00092955 | The frequency of TAT in AS region sequence |
| 86 | asseqfrequencyTA C | 0.00071859 | The frequency of TAC in AS region sequence |
| 87 | asseqfrequencyTA G | 0.00100277 | The frequency of TAG in AS region sequence |
| 88 | asseqfrequencyTT | 0.00325989 | The frequency of TT in AS region sequence |
| 89 | asseqfrequencyTT A | 0.00210937 | The frequency of TTA in AS region sequence |
| 90 | asseqfrequencyTT T | 0.00232819 | The frequency of TTT in AS region sequence |
| 91 | asseqfrequencyTT C | 0.00114105 | The frequency of TTC in AS region sequence |
| 92 | asseqfrequencyTT G | 0.00223335 | The frequency of TTG in AS region sequence |
| 93 | asseqfrequencyTC | 0.00106129 | The frequency of TC in AS region sequence |
| 94 | asseqfrequencyTC A | 0.00068795 | The frequency of TCA in AS region sequence |
| 95 | asseqfrequencyTC T | 0.00123544 | The frequency of TCT in AS region sequence |
| 96 | asseqfrequencyTC C | 0.00077003 | The frequency of TCC in AS region sequence |
| 97 | asseqfrequencyTC | 0.00078591 | The frequency of TCG in AS region |

| | | | |
|---|---|---|---|
| | G | | sequence |
| 98 | asseqfrequencyTG | 0.00172659 | The frequency of TG in AS region sequence |
| 99 | asseqfrequencyTGA | 0.00067501 | The frequency of TGA in AS region sequence |
| 100 | asseqfrequencyTGT | 0.00217143 | The frequency of TGT in AS region sequence |
| 101 | asseqfrequencyTGC | 0.00131095 | The frequency of TGC in AS region sequence |
| 102 | asseqfrequencyTGG | 0.00078633 | The frequency of TGG in AS region sequence |
| 103 | asseqfrequencyC | 0.00094733 | The frequency of C in AS region sequence |
| 104 | asseqfrequencyCA | 0.00177709 | The frequency of CA in AS region sequence |
| 105 | asseqfrequencyCAA | 0.00201401 | The frequency of CAA in AS region sequence |
| 106 | asseqfrequencyCAT | 0.00066235 | The frequency of CAT in AS region sequence |
| 107 | asseqfrequencyCAG | 0.00379374 | The frequency of CAG in AS region sequence |
| 108 | asseqfrequencyCT | 0.00460841 | The frequency of CT in AS region sequence |
| 109 | asseqfrequencyCTA | 0.00052012 | The frequency of CTA in AS region sequence |
| 110 | asseqfrequencyCTT | 0.00092103 | The frequency of CTT in AS region sequence |
| 111 | asseqfrequencyCTC | 0.00079791 | The frequency of CTC in AS region sequence |
| 112 | asseqfrequencyCTG | 0.00055332 | The frequency of CTG in AS region sequence |

| | | | |
|---|---|---|---|
| 113 | asseqfrequencyCC | 0.00059355 | The frequency of CC in AS region sequence |
| 114 | asseqfrequencyCCA | 0.00054019 | The frequency of CCA in AS region sequence |
| 115 | asseqfrequencyCCT | 0.00051827 | The frequency of CCT in AS region sequence |
| 116 | asseqfrequencyCG | 0.00099069 | The frequency of CG in AS region sequence |
| 117 | asseqfrequencyCGA | 0.00087573 | The frequency of CGA in AS region sequence |
| 118 | asseqfrequencyG | 0.0013486 | The frequency of G in AS region sequence |
| 119 | asseqfrequencyGA | 0.00160472 | The frequency of GA in AS region sequence |
| 120 | asseqfrequencyGAA | 0.00073121 | The frequency of GAA in AS region sequence |
| 121 | asseqfrequencyGAT | 0.00075458 | The frequency of GAT in AS region sequence |
| 122 | asseqfrequencyGAC | 0.00051875 | The frequency of GAC in AS region sequence |
| 123 | asseqfrequencyGAG | 0.00086106 | The frequency of GAG in AS region sequence |
| 124 | asseqfrequencyGT | 0.00295507 | The frequency of GT in AS region sequence |
| 125 | asseqfrequencyGTA | 0.00190334 | The frequency of GTA in AS region sequence |
| 126 | asseqfrequencyGTT | 0.00073332 | The frequency of GTT in AS region sequence |
| 127 | asseqfrequencyGTC | 0.00063169 | The frequency of GTC in AS region sequence |
| 128 | asseqfrequencyGT | 0.00068685 | The frequency of GTG in AS region |

| | | | |
|---|---|---|---|
| | G | | sequence |
| 129 | asseqfrequencyGC | 0.00111047 | The frequency of GC in AS region sequence |
| 130 | asseqfrequencyGCA | 0.00112206 | The frequency of GCA in AS region sequence |
| 131 | asseqfrequencyGCT | 0.00070271 | The frequency of GCT in AS region sequence |
| 132 | asseqfrequencyGG | 0.00135013 | The frequency of GG in AS region sequence |
| 133 | asseqfrequencyGGA | 0.00124547 | The frequency of GGA in AS region sequence |
| 134 | asseqfrequencyGGT | 0.00070964 | The frequency of GGT in AS region sequence |
| 135 | asseqfrequencyGGC | 0.0006017 | The frequency of GGC in AS region sequence |
| 136 | asseqfrequencyGGG | 0.00068444 | The frequency of GGG in AS region sequence |
| 137 | asseqdistr1%A | 0.00726205 | The distribution(position/length) of 1% A in AS region sequence |
| 138 | asseqdistr25%A | 0.00679882 | The distribution(position/length) of 25% A in AS region sequence |
| 139 | asseqdistr50%A | 0.00161645 | The distribution(position/length) of 50% A in AS region sequence |
| 140 | asseqdistr75%A | 0.00142922 | The distribution(position/length) of 75% A in AS region sequence |
| 141 | asseqdistr100%A | 0.0075252 | The distribution(position/length) of 100% A in AS region sequence |
| 142 | asseqdistr1%T | 0.00601582 | The distribution(position/length) of 1% T in AS region sequence |
| 143 | asseqdistr25%T | 0.0024819 | The distribution(position/length) of 25% T in AS region sequence |

| | | | |
|---|---|---|---|
| 144 | asseqdistr50%T | 0.00409853 | The distribution(position/length) of 50% T in AS region sequence |
| 145 | asseqdistr75%T | 0.004453 | The distribution(position/length) of 75% T in AS region sequence |
| 146 | asseqdistr100%T | 0.00978825 | The distribution(position/length) of 100% T in AS region sequence |
| 147 | asseqdistr1%C | 0.00352705 | The distribution(position/length) of 1% C in AS region sequence |
| 148 | asseqdistr25%C | 0.0019445 | The distribution(position/length) of 25% C in AS region sequence |
| 149 | asseqdistr50%C | 0.00090629 | The distribution(position/length) of 50% C in AS region sequence |
| 150 | asseqdistr75%C | 0.00240864 | The distribution(position/length) of 75% C in AS region sequence |
| 151 | asseqdistr100%C | 0.00857147 | The distribution(position/length) of 100% C in AS region sequence |
| 152 | asseqdistr1%G | 0.01593786 | The distribution(position/length) of 1% G in AS region sequence |
| 153 | asseqdistr25%G | 0.01090831 | The distribution(position/length) of 25% G in AS region sequence |
| 154 | asseqdistr50%G | 0.00478994 | The distribution(position/length) of 50% G in AS region sequence |
| 155 | asseqdistr75%G | 0.00422594 | The distribution(position/length) of 75% G in AS region sequence |
| 156 | asseqdistr100%G | 0.00723644 | The distribution(position/length) of 100% G in AS region sequence |
| 157 | asseqdonerGT | 0.00131268 | Is there GT in doner of AS region sequence (1 for yes, 0 for no) |
| 158 | asseqacceptorGT | 0.02884584 | Is there GT in acceptor of AS region sequence (1 for yes, 0 for no) |
| 159 | asseqdonerGC | 0.00222911 | Is there GC in doner of AS region |

| | | | |
|---|---|---|---|
| | | | sequence (1 for yes, 0 for no) |
| 160 | asseqacceptorGC | 0.00105832 | Is there GC in acceptor of AS region sequence (1 for yes, 0 for no) |
| 161 | asseqdonerAT | 0.00205132 | Is there AT in doner of AS region sequence (1 for yes, 0 for no) |
| 162 | asseqacceptorAT | 0.00089301 | Is there AT in acceptor of AS region sequence (1 for yes, 0 for no) |
| 163 | asseqdonerAG | 0.0279886 | Is there AG in doner of AS region sequence (1 for yes, 0 for no) |
| 164 | asseqacceptorAG | 0.00086802 | Is there AG in acceptor of AS region sequence (1 for yes, 0 for no) |
| 165 | upfrequencyTC | 0.00054334 | The frequency of TC in upstream sequence |
| 166 | upfrequencyC | 0.00057141 | The frequency of C in upstream sequence |
| 167 | updistr100%G | 0.00087752 | The distribution(position/length) of 100% G in upstream sequence |
| 168 | updonerGT | 0.00056503 | Is there GT in doner of upstream sequence (1 for yes, 0 for no) |
| 169 | updonerAG | 0.0010452 | Is there AG in doner of upstream sequence (1 for yes, 0 for no) |
| 170 | downfrequencyTC | 0.00063755 | The frequency of TC in downstream sequence |
| 171 | downfrequencyTCT | 0.00057412 | The frequency of TCT in downstream sequence |
| 172 | downfrequencyCT | 0.00065043 | The frequency of CT in downstream sequence |
| 173 | downdistr100%G | 0.00058024 | The distribution(position/length) of 100% G in downstream sequence |
| 174 | downdonerAG | 0.00055697 | Is there AG in doner of in downstream sequence (1 for yes, 0 for no) |

| | | | |
|---|---|---|---|
| 175 | up30as30if%3 | 0.00526169 | Whethere the length of upstream30 + AS 30bp sequence divisible by three |
| 176 | up30as30GC | 0.00102788 | The GC contant of upstream30AS 30bp + AS 30bp sequence |
| 177 | up30as30numberTAA | 0.00062303 | The number of stopdocon TAA in upstream30AS 30bp + AS 30bp sequence |
| 178 | up30as30frequencyAG | 0.00083881 | The frequency of AG in upstream30AS 30bp + AS 30bp sequence |
| 179 | up30as30frequencyAGG | 0.00063295 | The frequency of AGG in upstream30AS 30bp + AS 30bp sequence |
| 180 | up30as30frequencyTA | 0.00071584 | The frequency of TA in upstream30AS 30bp + AS 30bp sequence |
| 181 | up30as30frequencyTAA | 0.00066461 | The frequency of TAA in upstream30AS 30bp + AS 30bp sequence |
| 182 | up30as30frequencyTTT | 0.00054375 | The frequency of TTT in upstream30AS 30bp + AS 30bp sequence |
| 183 | up30as30frequencyCAG | 0.00129916 | The frequency of CAG in upstream30AS 30bp + AS 30bp sequence |
| 184 | up30as30frequencyG | 0.00057868 | The frequency of G in upstream30AS 30bp + AS 30bp sequence |
| 185 | up30as30frequencyGA | 0.00090161 | The frequency of GA in upstream30AS 30bp + AS 30bp sequence |
| 186 | up30as30frequencyGAT | 0.0005604 | The frequency of GAT in upstream30AS 30bp + AS 30bp sequence |
| 187 | up30as30frequencyGTA | 0.00224494 | The frequency of GTA in upstream30AS 30bp + AS 30bp sequence |
| 188 | up30as30frequencyGGA | 0.00056531 | The frequency of GGA in upstream30AS 30bp + AS 30bp sequence |
| 189 | up30as30frequencyGGT | 0.00241256 | The frequency of GGT in upstream30AS 30bp + AS 30bp sequence |

| | | | |
|---|---|---|---|
| 190 | up30as30frequencyGGC | 0.00053686 | The frequency of GGC in upstream30AS 30bp + AS 30bp sequence |
| 191 | up30as30distr100%A | 0.00146039 | The distribution(position/length) of 100% A in upstream30AS 30bp + AS 30bp sequence |
| 192 | up30as30distr50%T | 0.00124887 | The distribution(position/length) of 50% T in upstream30AS 30bp + AS 30bp sequence |
| 193 | up30as30distr75%T | 0.00116163 | The distribution(position/length) of 75% T in upstream30AS 30bp + AS 30bp sequence |
| 194 | up30as30distr100%T | 0.00142157 | The distribution(position/length) of 100% T in upstream30AS 30bp + AS 30bp sequence |
| 195 | up30as30distr50%C | 0.00056688 | The distribution(position/length) of 50% C in upstream30AS 30bp + AS 30bp sequence |
| 196 | up30as30distr75%C | 0.00078279 | The distribution(position/length) of 75% C in upstream30AS 30bp + AS 30bp sequence |
| 197 | up30as30distr25%G | 0.00052417 | The distribution(position/length) of 25% G in upstream30AS 30bp + AS 30bp sequence |
| 198 | up30as30distr50%G | 0.00143519 | The distribution(position/length) of 50% G in upstream30AS 30bp + AS 30bp sequence |
| 199 | up30as30distr75%G | 0.00368218 | The distribution(position/length) of 75% G in upstream30AS 30bp + AS 30bp sequence |
| 200 | up30as30distr100%G | 0.00300135 | The distribution(position/length) of 100% G in upstream30AS 30bp + AS 30bp sequence |
| 201 | up30as30donerGT | 0.00075511 | Is there GT in doner of upstream30AS |

| | | | |
|---|---|---|---|
| | | | 30bp + AS 30bp sequence (1 for yes, 0 for no) |
| 202 | up30as30donerGC | 0.00082364 | Is there GC in doner of upstream30AS 30bp + AS 30bp sequence (1 for yes, 0 for no) |
| 203 | up30as30donerAT | 0.00057632 | Is there AT in doner of upstream30AS 30bp + AS 30bp sequence (1 for yes, 0 for no) |
| 204 | up30as30donerAG | 0.00882961 | Is there AG in doner of upstream30AS 30bp + AS 30bp sequence (1 for yes, 0 for no) |
| 205 | up30down30distr50%G | 0.00066765 | The distribution(position/length) of 50% G in upstream 30 + downstream 30 sequence |
| 206 | up30down30distr75%G | 0.00055362 | The distribution(position/length) of 75% G in upstream 30 + downstream 30 sequence |
| 207 | as30down30if%3 | 0.00412092 | Whethere the length of AS 30bp + downstream 30bp sequence divisible by three |
| 208 | as30down30GC | 0.00102468 | The GC contant of AS 30bp + downstream 30bp sequence |
| 209 | as30down30frequencyAAG | 0.00055681 | The frequency of AAG in AS 30bp + downstream 30bp sequence |
| 210 | as30down30frequencyAG | 0.00066375 | The frequency of AG in AS 30bp + downstream 30bp sequence |
| 211 | as30down30frequencyAGG | 0.00057326 | The frequency of AGG in AS 30bp + downstream 30bp sequence |
| 212 | as30down30frequencyT | 0.00060534 | The frequency of T in AS 30bp + downstream 30bp sequence |
| 213 | as30down30frequencyTTT | 0.00064802 | The frequency of TTT in AS 30bp + downstream 30bp sequence |

| | | | |
|---|---|---|---|
| 214 | as30down30frequencyTC | 0.00051054 | The frequency of TC in AS 30bp + downstream 30bp sequence |
| 215 | as30down30frequencyTG | 0.00051856 | The frequency of TG in AS 30bp + downstream 30bp sequence |
| 216 | as30down30frequencyC | 0.00052519 | The frequency of C in AS 30bp + downstream 30bp sequence |
| 217 | as30down30frequencyCAG | 0.00161718 | The frequency of CAG in AS 30bp + downstream 30bp sequence |
| 218 | as30down30frequencyGAG | 0.00060058 | The frequency of GAG in AS 30bp + downstream 30bp sequence |
| 219 | as30down30distr1%A | 0.00071615 | The distribution(position/length) of 1% A in AS 30bp + downstream 30bp sequence |
| 220 | as30down30distr1%T | 0.00065785 | The distribution(position/length) of 1% T in AS 30bp + downstream 30bp sequence |
| 221 | as30down30distr25%T | 0.00051918 | The distribution(position/length) of 25% T in AS 30bp + downstream 30bp sequence |
| 222 | as30down30distr50%T | 0.00121401 | The distribution(position/length) of 50% T in AS 30bp + downstream 30bp sequence |
| 223 | as30down30distr75%T | 0.00051052 | The distribution(position/length) of 75% T in AS 30bp + downstream 30bp sequence |
| 224 | as30down30distr1%G | 0.00161531 | The distribution(position/length) of 1% G in AS 30bp + downstream 30bp sequence |
| 225 | as30down30distr25%G | 0.00073252 | The distribution(position/length) of 25% G in AS 30bp + downstream 30bp sequence |
| 226 | as30down30distr5 | 0.00137949 | The distribution(position/length) of 50% G in AS 30bp + downstream 30bp |

| | | | |
|---|---|---|---|
| | 0%G | | sequence |
| 227 | as30down30distr75%G | 0.00054911 | The distribution(position/length) of 75% G in AS 30bp + downstream 30bp sequence |
| 228 | as30down30acceptorGT | 0.00237995 | Is there GT in acceptor of AS 30bp + downstream 30bp sequence (1 for yes, 0 for no) |
| 229 | as30down30acceptorAG | 0.00103183 | Is there AG in acceptor of AS 30bp + downstream 30bp sequence (1 for yes, 0 for no) |
| 230 | up50as50if%3 | 0.00313826 | Whethere the length of upstream 50bp + AS 50bp sequence divisible by three |
| 231 | up50as50GC | 0.00248468 | The GC contant of upstream 50bp + AS 50bp sequence |
| 232 | up50as50numberTAA | 0.00090647 | The numberof stopdocon TAA in upstream 50bp + AS 50bp sequence |
| 233 | up50as50frequencyAG | 0.00066188 | The frequency of AG in upstream 50bp + AS 50bp sequence |
| 234 | up50as50frequencyAGG | 0.00052156 | The frequency of AGG in upstream 50bp + AS 50bp sequence |
| 235 | up50as50frequencyT | 0.00055684 | The frequency of T in upstream 50bp + AS 50bp sequence |
| 236 | up50as50frequencyTA | 0.00076884 | The frequency of TA in upstream 50bp + AS 50bp sequence |
| 237 | up50as50frequencyTAA | 0.00107979 | The frequency of TAA in upstream 50bp + AS 50bp sequence |
| 238 | up50as50frequencyTTA | 0.00061682 | The frequency of TTA in upstream 50bp + AS 50bp sequence |
| 239 | up50as50frequencyCAG | 0.00110433 | The frequency of CAG in upstream 50bp + AS 50bp sequence |
| 240 | up50as50frequenc | 0.00076794 | The frequency of G in upstream 50bp + |

| | | | |
|---|---|---|---|
| | yG | | AS 50bp sequence |
| 241 | up50as50frequencyGA | 0.00070365 | The frequency of GA in upstream 50bp + AS 50bp sequence |
| 242 | up50as50frequencyGAG | 0.00051534 | The frequency of GAG in upstream 50bp + AS 50bp sequence |
| 243 | up50as50frequencyGTA | 0.00156303 | The frequency of GTA in upstream 50bp + AS 50bp sequence |
| 244 | up50as50frequencyGGA | 0.00060143 | The frequency of GGA in upstream 50bp + AS 50bp sequence |
| 245 | up50as50frequencyGGT | 0.00100631 | The frequency of GGT in upstream 50bp + AS 50bp sequence |
| 246 | up50as50distr100%A | 0.0013585 | The distribution(position/length) of 100% A in upstream 50bp + AS 50bp sequence |
| 247 | up50as50distr50%T | 0.00222866 | The distribution(position/length) of 50% T in upstream 50bp + AS 50bp sequence |
| 248 | up50as50distr75%T | 0.00060388 | The distribution(position/length) of 75% T in upstream 50bp + AS 50bp sequence |
| 249 | up50as50distr100%T | 0.00220456 | The distribution(position/length) of 100% T in upstream 50bp + AS 50bp sequence |
| 250 | up50as50distr50%C | 0.00052123 | The distribution(position/length) of 50% C in upstream 50bp + AS 50bp sequence |
| 251 | up50as50distr75%C | 0.00059727 | The distribution(position/length) of 75% C in upstream 50bp + AS 50bp sequence |
| 252 | up50as50distr100%C | 0.00066252 | The distribution(position/length) of 100% C in upstream 50bp + AS 50bp sequence |

| | | | |
|---|---|---|---|
| 253 | up50as50distr1%G | 0.0005147 | The distribution(position/length) of 1% G in upstream 50bp + AS 50bp sequence |
| 254 | up50as50distr25%G | 0.00066241 | The distribution(position/length) of 25% G in upstream 50bp + AS 50bp sequence |
| 255 | up50as50distr50%G | 0.00175489 | The distribution(position/length) of 50% G in upstream 50bp + AS 50bp sequence |
| 256 | up50as50distr75%G | 0.00172491 | The distribution(position/length) of 75% G in upstream 50bp + AS 50bp sequence |
| 257 | up50as50distr100%G | 0.00265711 | The distribution(position/length) of 100% G in upstream 50bp + AS 50bp sequence |
| 258 | up50as50donerGT | 0.00080078 | Is there GT in doner of upstream 50bp + AS 50bp sequence (1 for yes, 0 for no) |
| 259 | up50as50donerGC | 0.00072178 | Is there GC in doner of upstream 50bp + AS 50bp sequence (1 for yes, 0 for no) |
| 260 | up50as50donerAT | 0.00075531 | Is there AT in doner of upstream 50bp + AS 50bp sequence (1 for yes, 0 for no) |
| 261 | up50as50donerAG | 0.00973815 | Is there AG in doner of upstream 50bp + AS 50bp sequence (1 for yes, 0 for no) |
| 262 | up50down50distr25%C | 0.00056377 | The distribution(position/length) of 25% C in upstream 50bp + downstream 50bp sequence (1 for yes, 0 for no) |
| 263 | up50down50distr50%C | 0.00064309 | The distribution(position/length) of 50% C in upstream 50bp + downstream 50bp sequence (1 for yes, 0 for no) |
| 264 | up50down50distr50%G | 0.00066223 | The distribution(position/length) of 50% G in upstream 50bp + downstream 50bp sequence (1 for yes, 0 for no) |
| 265 | up50down50distr7 | 0.00055767 | The distribution(position/length) of 75% G in upstream 50bp + downstream |

| | | | |
|---|---|---|---|
| | 5%G | | 50bp sequence (1 for yes, 0 for no) |
| 266 | up50down50distr100%G | 0.00054658 | The distribution(position/length) of 100% G in upstream 50bp + downstream 50bp sequence (1 for yes, 0 for no) |
| 267 | as50down50if%3 | 0.00369669 | Whethere the length of AS 50bp + downstream 50bp sequence divisible by three |
| 268 | as50down50GC | 0.00349382 | The GC contant of GC in AS 50bp + downstream 50bp sequence |
| 269 | as50down50numberTAA | 0.0008039 | The number of stopdocon TAA in AS 50bp + downstream 50bp sequence |
| 270 | as50down50frequencyAAG | 0.00053626 | The frequency of AAG in AS 50bp + downstream 50bp sequence |
| 271 | as50down50frequencyAG | 0.000601 | The frequency of AG in AS 50bp + downstream 50bp sequence |
| 272 | as50down50frequencyAGA | 0.00051576 | The frequency of AGA in AS 50bp + downstream 50bp sequence |
| 273 | as50down50frequencyAGG | 0.00060834 | The frequency of AGG in AS 50bp + downstream 50bp sequence |
| 274 | as50down50frequencyT | 0.00096353 | The frequency of T in AS 50bp + downstream 50bp sequence |
| 275 | as50down50frequencyTAA | 0.00052375 | The frequency of TAA in AS 50bp + downstream 50bp sequence |
| 276 | as50down50frequencyTT | 0.00064601 | The frequency of TT in AS 50bp + downstream 50bp sequence |
| 277 | as50down50frequencyTTA | 0.00061379 | The frequency of TTA in AS 50bp + downstream 50bp sequence |
| 278 | as50down50frequencyTC | 0.00060552 | The frequency of TC in AS 50bp + downstream 50bp sequence |
| 279 | as50down50frequencyTCG | 0.0005862 | The frequency of TCG in AS 50bp + downstream 50bp sequence |

| | | | |
|---|---|---|---|
| 280 | as50down50frequencyTGT | 0.0007312 | The frequency of TGT in AS 50bp + downstream 50bp sequence |
| 281 | as50down50frequencyC | 0.00053764 | The frequency of C in AS 50bp + downstream 50bp sequence |
| 282 | as50down50frequencyCAG | 0.00073511 | The frequency of CAG in AS 50bp + downstream 50bp sequence |
| 283 | as50down50frequencyCTC | 0.00056771 | The frequency of CTC in AS 50bp + downstream 50bp sequence |
| 284 | as50down50frequencyCG | 0.00056719 | The frequency of CG in AS 50bp + downstream 50bp sequence |
| 285 | as50down50frequencyCGA | 0.00051527 | The frequency of CGA in AS 50bp + downstream 50bp sequence |
| 286 | as50down50frequencyG | 0.00068001 | The frequency of G in AS 50bp + downstream 50bp sequence |
| 287 | as50down50distr1%A | 0.0008114 | The distribution(position/length) of 1% A in AS 50bp + downstream 50bp sequence |
| 288 | as50down50distr1%T | 0.00154536 | The distribution(position/length) of 1% T in AS 50bp + downstream 50bp sequence |
| 289 | as50down50distr50%T | 0.00121784 | The distribution(position/length) of 50% T in AS 50bp + downstream 50bp sequence |
| 290 | as50down50distr1%C | 0.00053517 | The distribution(position/length) of 1% C in AS 50bp + downstream 50bp sequence |
| 291 | as50down50distr1%G | 0.00166606 | The distribution(position/length) of 1% G in AS 50bp + downstream 50bp sequence |
| 292 | as50down50distr25%G | 0.00117026 | The distribution(position/length) of 25% G in AS 50bp + downstream 50bp sequence |

| | | | |
|---|---|---|---|
| 293 | as50down50distr50%G | 0.00155005 | The distribution(position/length) of 50% G in AS 50bp + downstream 50bp sequence |
| 294 | as50down50distr75%G | 0.00051549 | The distribution(position/length) of 75% G in AS 50bp + downstream 50bp sequence |
| 295 | as50down50distr100%G | 0.00052391 | The distribution(position/length) of 100% G in AS 50bp + downstream 50bp sequence |
| 296 | as50down50acceptorGT | 0.00348638 | Is there GT in acceptor of AS 50bp + downstream 50bp sequence (1 for yes, 0 for no) |
| 297 | as50down50acceptorAG | 0.00065119 | Is there AG in acceptor of AS 50bp + downstream 50bp sequence (1 for yes, 0 for no) |
